# Sequential deactivation across the thalamus-hippocampus-mPFC pathway during loss of consciousness

**DOI:** 10.1101/2024.05.20.594986

**Authors:** Xiaoai Chen, Samuel R. Cramer, Dennis C.Y. Chan, Xu Han, Nanyin Zhang

**Author notes:** **Corresponding author:** Nanyin Zhang, Ph.D. –.

## Abstract

How consciousness is lost in states such as sleep or anesthesia remains a mystery. To gain insight into this phenomenon, we conducted concurrent recordings of electrophysiology signals in the anterior cingulate cortex and whole-brain functional magnetic resonance imaging (fMRI) in rats exposed to graded propofol, undergoing the transition from consciousness to unconsciousness. Our results reveal that upon the loss of consciousness (LOC), as indicated by the loss of righting reflex, there is a sharp increase in low-frequency power of the electrophysiological signal. Additionally, simultaneously measured fMRI signals exhibit a cascade of deactivation across a pathway including the hippocampus, thalamus, and medial prefrontal cortex (mPFC) surrounding the moment of LOC, followed by a broader increase in brain activity across the cortex during sustained unconsciousness. Furthermore, sliding window analysis demonstrates a temporary increase in synchrony of fMRI signals across the hippocampus-thalamus-mPFC pathway preceding LOC. These data suggest that LOC might be triggered by sequential activities in the hippocampus, thalamus and mPFC, while wide-spread activity increases in other cortical regions commonly observed during anesthesia-induced unconsciousness might be a consequence, rather than a cause of LOC. Taken together, our study identifies a cascade of neural events unfolding as the brain transitions into unconsciousness, offering critical insight into the systems-level neural mechanisms underpinning LOC.

## Introduction

The phenomenon of how consciousness is lost in states such as sleep or anesthesia remains a subject of mystery. Investigating the loss of consciousness (LOC) induced by general anesthesia provides a promising avenue for research due to the robustness of experimental control it offers and its pivotal role in medical care. Anesthesia-induced unconsciousness (AIU) typically begins with the administration of a fast-acting anesthetic agent, quickly leading to LOC, a state that can be modulated and/or maintained by varying the dose of anesthetics^1^. While the molecular and cellular effects of different anesthetics are generally well-understood, the neural mechanism of AIU at the systems level, especially during the transition of LOC, remains elusive.

Neuroimaging methods play a crucial role in unraveling the brain-wide neural activity changes associated with AIU. At the regional level, studies have revealed alterations in local brain activity under propofol-induced anesthesia across multiple brain regions, including anterior frontal regions, the temporal pole, hippocampus, and amygdala^2,3^. Moving up in scale, analyses of brain networks have demonstrated significant changes in cortico-cortical connectivity, particularly within the default mode network (DMN) and frontoparietal network^4,5,6^. Additionally, reports have indicated that thalamic-cortical connectivity, particularly the connectivity between the nonspecific thalamus and cortex, is significantly different during AIU^7,8,9^. However, while these findings are often observed in resting-state functional magnetic resonance imaging (rsfMRI) studies, their mechanistic relationship to unconsciousness remains unclear. Moreover, most rsfMRI research has focused on the stable state of general anesthesia, potentially overlooking dynamic brain changes during the transition to LOC.

LOC can occur rapidly, within seconds to minutes, and is associated with several characteristic neurophysiological changes^1^. For instance, slow fluctuations in cortical electrophysiological signals were found to rapidly increase immediately after LOC and became phase-coupled with neural spiking and alpha band amplitude^10,11^. Cortico-cortical and thalamocortical synchronization in the alpha band of the electrophysiological signal were also reported during LOC^12,13^. However, due to the limited spatial coverage of electrophysiology, understanding global brain changes during LOC remains challenging. Meanwhile, rsfMRI alone, limited by its temporal resolution due to the slow hemodynamic response^14^, cannot track fast-changing neural dynamics across the brain effectively. Therefore, simultaneous electrophysiology-rsfMRI recording presents a promising approach to investigate whole-brain spatiotemporal dynamics during LOC, offering high temporal resolution from electrophysiology and high spatial resolution and whole-brain coverage from fMRI.

In this study, we conducted simultaneous electrophysiology-rsfMRI recordings in graded propofol-exposed rats undergoing the transition from consciousness to unconsciousness. Propofol was selected for its widespread use in medical procedures^15^. Electrophysiological signals were recorded from the left anterior cingulate cortex (ACC), a region indicative of the level of consciousness^16^. Additionally, the loss of righting reflex (LORR) test was conducted outside of the MRI scanner to provide a behavioral assessment of animals’ consciousness levels during the anesthesia paradigm. Our results demonstrate a sharp increase in low-frequency power in the electrophysiological signal, aligning well with the timing of LORR. Moreover, based on this electrophysiology signature-informed LOC, we observed a cascade of deactivation in the blood-oxygen-level-dependent (BOLD) intensity across a pathway including the hippocampus, thalamus, and medial prefrontal cortex (mPFC) around the moment of LOC, followed by a broader increase in BOLD signal across the cortex. Furthermore, sliding window analysis revealed a transient increase in global synchrony preceding LOC, particularly across the hippocampus-thalamus-mPFC pathway. Collectively, our study identifies a cascade of neural events unfolding as the brain transitions into unconsciousness.

## Results

To explore the spatiotemporal dynamics of brain activity during the LOC, we conducted simultaneous recordings of electrophysiological signals in the anterior cingulate cortex (ACC) and whole-brain rsfMRI in rats subjected to graded propofol administration (Fig. 1a; low dose: 20 mg/kg/h; high dose: 40 mg/kg/h). The ACC electrode was implanted chronically, and its placement was confirmed using T2-weighted structural MRI images (Fig. S1). We employed a template regression approach^17,18^ to remove magnetic resonance artifacts from the raw electrophysiology signal, yielding denoised electrophysiological data (Fig. S2). Subsequently, the preprocessed electrophysiological signal was band-pass filtered (0.2 - 100 Hz) to obtain the local field potential (LFP), from which band-specific LFP power was derived using conventional frequency bands: slow wave (0.2-1 Hz), delta (1–4 Hz), theta (5–7 Hz), alpha (8–12 Hz), beta (13–30 Hz), and gamma (40–100 Hz). Fig. 1b depicts the cross correlations between LFP powers and the BOLD signal in the ACC across the LFP spectrum (top; band interval = 0.186 Hz) and for individual LFP bands (bottom) at the low dose of propofol. Notably, the local BOLD signal exhibits a negative correlation with low-frequency LFP powers but a positive correlation with high-frequency LFP powers, in line with our previous findings^17,18^. The lag of the BOLD signal is consistently around 1-3 seconds for all bands, aligning with the delay in the hemodynamic response function (HRF) previously reported in rodents^19,20,21^.

**Figure 1.**
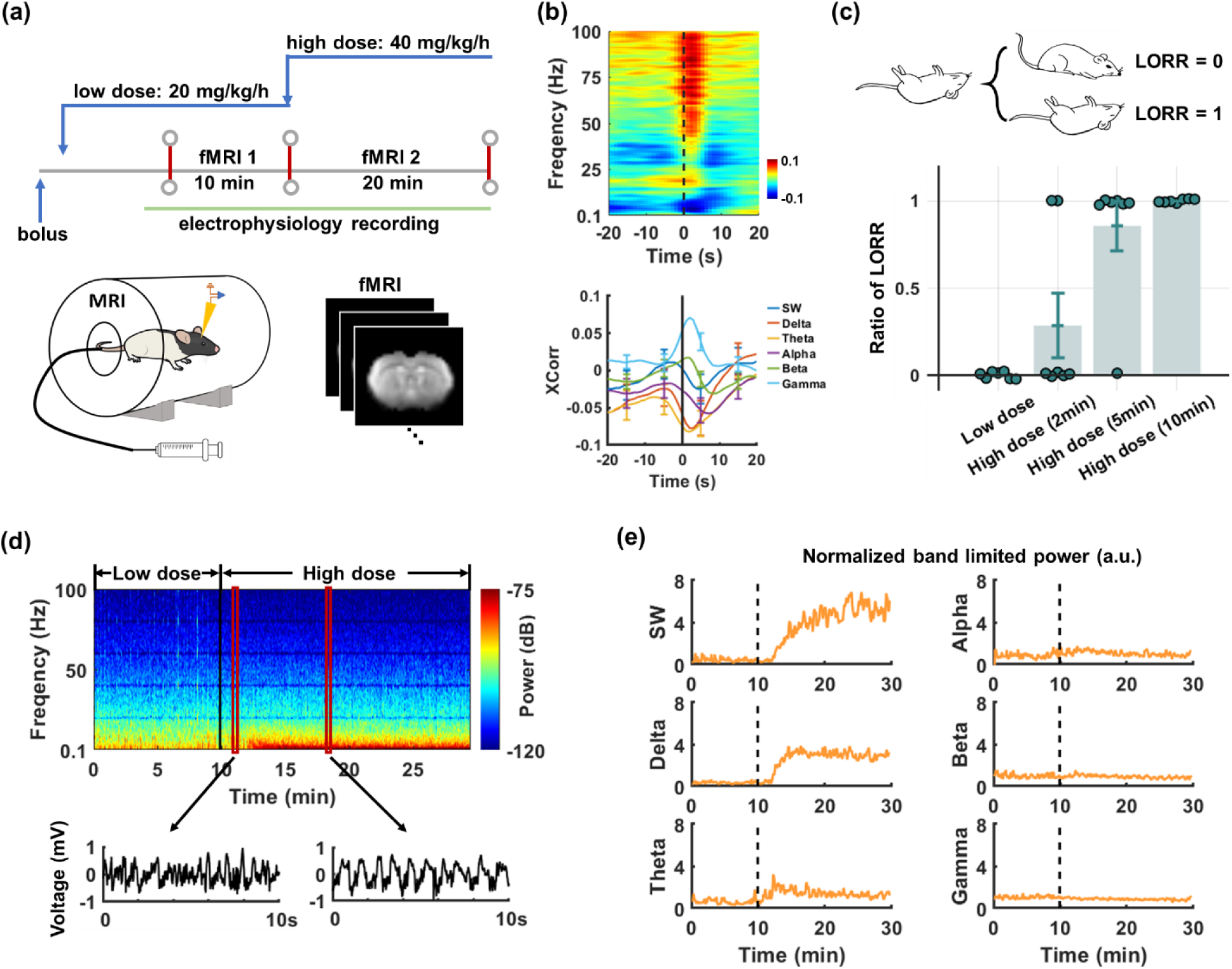
Simultaneous recordings of electrophysiology and fMRI signals during incremental doses of propofol. (**a**) Experimental design: Initially, a bolus of propofol (20 mg/kg) was injected, followed by the infusion of a low dose of propofol (20 mg/kg/h) for at least 40 min to achieve a steady state before the first rsfMRI scan (10 min) started. Subsequently, the propofol infusion rate was elevated to 40 mg/kg/h for 20 min, as the second rsfMRI scan was continuously acquired. (**b**) BOLD-electrophysiology cross correlations in the ACC at the low dose of propofol across the LFP spectrum (top, with the 0.186-Hz band interval) and for individual LFP bands (bottom). (**c**) LORR observed at different stages/times of propofol infusion. (**d**) Top: electrophysiology power spectrum of a representative scan. Bottom: LFP signals from selected windows (marked by red rectangles) within the exemplar power spectrum. (**e**) Normalized band limited power of the scan.

### Animals transition into a state of unconsciousness within 10 min following the initiation of high-dose propofol administration

To evaluate the level of consciousness under our anesthesia paradigm, we conducted a Loss of Righting Reflex (LORR) test^22^ outside of the scanner, employing the same infusion protocol as utilized during scanning (see Fig. 1a). Each animal underwent LORR testing at the low dose and at 2, 5, and 10 min following the initiation of high-dose propofol infusion, with the rat swiftly placed in a supine position. A complete LORR was recorded if the rat failed to return to the prone position within 30 s ^23^. Notably, no rats exhibited LORR at the low dose of propofol. However, 28.6% of rats lost their righting reflex within 2 min following the initiation of high-dose propofol administration. This percentage increased to 85.7% at 5 min and reached 100% at 10 min post-infusion (see Fig. 1c). These findings suggest that the LOC occurs during the transition from low to high dose of propofol, specifically within the first 10 min following the onset of high-dose infusion.

### Low-frequency LFP powers exhibited a significant and robust increase following the initiation of high-dose propofol administration

Given that the LOC occurs within the first 10 min following the onset of high-dose infusion, we specifically examined the electrophysiological signal during this period. Across animals, following the infusion of high-dose propofol a substantial increase was observed in average powers across multiple frequency bands including slow wave (0.2-1Hz), delta (1-4 Hz), theta (4-8 Hz), alpha (8-12 Hz), and beta (13-30 Hz) bands (Figs. 1d-e). Notably, the most prominent changes were observed in the slow-wave and delta bands, while alterations in higher-frequency bands (gamma (40–100 Hz), beta, alpha, and theta) were comparatively more modest (Fig. 1e). In addition, the power increase in these bands appeared to be abrupt and synchronized, particularly evident in the low-frequency bands (see Figs. 1d-e). As a result, the onset of LFP power increase was determined for each scan based on slow-wave and delta-band data. Aligning with this onset, for each LFP band we statistically examined the band-limited power amplitude for individual 10-s time bins within the interval from 20 s before to 800 s after the onset (Fig. 2a). Given that LFP powers continued to increase during this period, for each band we only reported the first time bin with a p-value < 0.05, as well as the percentage change of the band-limited power after reaching a plateau (∼300 s to 800 s post-onset) relative to the baseline (defined as the interval of −120 s to −10 s relative to the onset, Fig. 2b). Following the onset, a significant increase was observed in slow-wave power (p_(0∼10 s after LOC)_ = 7 × 10^-3^), reaching 485.5% ± 8.7% around 300 s after the onset. Similarly, delta band power exhibited a significant increase (p_(0∼10 s after LOC)_ = 1.3 × 10^-4^), rising to 570.9% ± 6.4% at the plateau. In comparison, changes in other frequency bands were relatively subtle but still significant: theta band, p_(0∼10 s after LOC)_ = 5.1 × 10^-5^, 296.6% ± 1.8% at the plateau; alpha band, p_(0∼10 s after LOC)_ = 2 × 10^-6^, 242.7% ± 1.9% at the plateau; and beta band, p-value_(0∼10 s after LOC)_ = 7.8× 10^-3^, 149.4%± 0.9% at the plateau. Additionally, gamma band power showed a significant but delayed increase from 160 s to 540 s after the onset. In summary, band-specific LFP powers began to increase after the initiation of high-dose propofol infusion and remained elevated throughout the remaining recording period, with the slow-wave and delta bands exhibiting the most pronounced changes.

**Figure 2.**
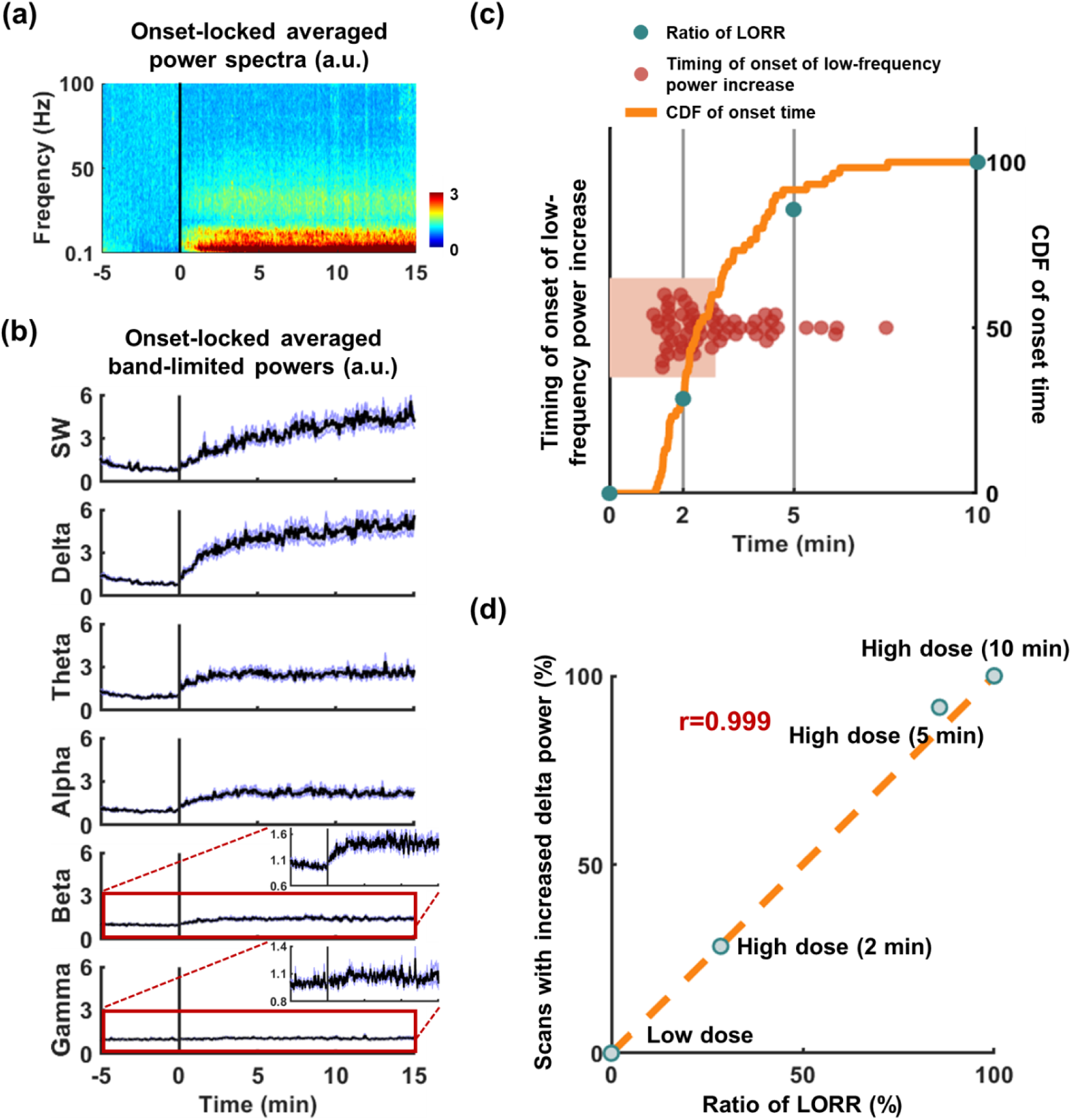
Simultaneous occurrence of low-frequency LFP power increase and loss of righting reflex (LORR). (**a**) Normalized power spectra, time-locked to the onset of slow-wave and delta-band power increase and averaged across scans. (**b**) Time-locked and averaged band-limited powers. (**c**) Distribution of timing following high-dose propofol infusion when the onset of low-frequency LFP power increase was observed, along with the portion of scans showing such increase at different times post-infusion, depicted using a cumulative distribution function (CDF). This portion was superimposed with the portion of animals exhibiting LORR at 0, 2, 5, and 10 min post-infusion. (**d**) Correlation between the percentage of animals exhibiting LORR at varying time points following high-dose infusion and the percentage of scans showing the onset of low-frequency LFP power increase within the corresponding time intervals.

### Onset of low-frequency LFP power increase aligned with behaviorally determined LOC

Both the LORR and the increase of low-frequency LFP powers were observed within 10 min following the initiation of high-dose propofol infusion. This prompted an inquiry into the temporal relationship between behaviorally determined LOC and the onset of LFP power increase. To explore this relationship, we compared the timing of the onset of low-frequency (slow wave and delta band) power increase with instances of LORR occurrence under separate stages/times of propofol infusion (see Figs. 2c-d). Notably, under the low dose of propofol, neither LORR nor low-frequency LFP power increase was evident across any of the animals. However, within 2 min post-commencement of high-dose propofol infusion, 28.3% rats manifested an abrupt increase of low-frequency LFP power, and 28.6% rats exhibited LORR. Within 5 min following high-dose administration, 91.7% rats displayed pronounced changes in low-frequency LPF power, correlating with LORR in 85.7% of cases. By the 10-min mark post-high-dose infusion, 100% rats exhibited a marked increase in low-frequency LFP power, coinciding with LORR in all rats (Fig. 2c). Remarkably, there was an exceptionally high correlation between the proportion of animals displaying LORR at various time points subsequent to high-dose infusion and the proportion of scans revealing low-frequency LFP power escalation during the same intervals (r = 0.999, Fig. 2d), indicating a robust connection between these two metrics. Consequently, this study posits that the escalation in low-frequency LFP power serves as a neurophysiological indicator of behaviorally determined LOC, consistent with existing literature findings^10,24,11,13,25^. This assertation led to the selection of the onset of low-frequency LFP power increase as an electrophysiology-based marker of LOC, allowing for further investigation of whole-brain alterations surrounding LOC using concurrently acquired rsfMRI data.

### As the LOC unfolds, a cascade of brain activities within the hippocampus-thalamus-medial prefrontal cortex pathway was observed

Using electrophysiology-informed LOC (hereafter referred to as ‘LOC’, defined as time zero), we systematically investigated the spatiotemporal dynamics of brain activity surrounding LOC. The BOLD intensity of each of the individual brain regions of interest (ROIs) during the time interval of −100 s to 300 s was normalized to baseline, defined as the averaged BOLD intensity of the ROI from −100 s to −45 s (Fig. 3a). Notably, both the thalamus and the medial prefrontal cortex (mPFC) exhibit a similar temporal pattern: a transient decrease in BOLD intensity near the time of LOC, followed by an increase and eventual plateau elevated above baseline post-LOC (Figs. 3b-c). Importantly, this pattern remains consistent across individual nuclei in the thalamus (Fig. S3) and individual subdivisions in the mPFC (Fig. S4).

**Figure 3.**
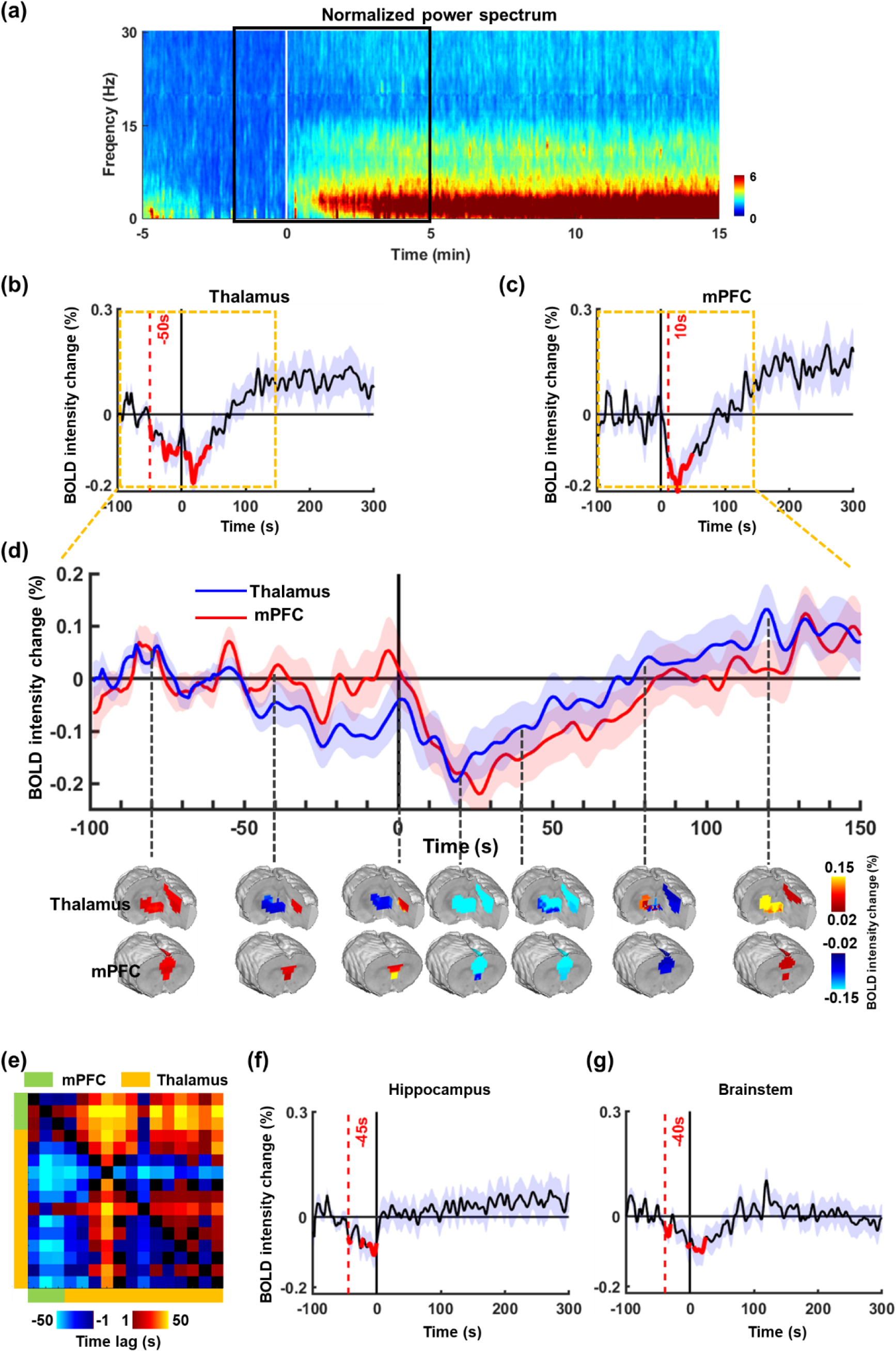
Change of BOLD signal intensity around electrophysiology-informed LOC. (**a**) The normalized power spectrum, time-locked to the electrophysiology-informed LOC and averaged across scans. The rsfMRI data were then segmented to a window from - 100 s to 300 s around the electrophysiology-informed LOC, as marked by the black box. (**b-d**) Both the thalamus and the medial prefrontal cortex (mPFC) exhibit a similar temporal pattern: a transient decrease in BOLD intensity near the time of LOC, followed by an increase and eventual plateau elevated above baseline post-LOC. (**b**) BOLD intensity change around LOC in the thalamus. (**c**) BOLD intensity change around LOC in the mPFC. The red curves represent time bins with significantly lower BOLD intensity (p-value < 0.05, two-tailed t-tests, uncorrected) than baseline (−100 s to −45 s before LOC) in each region. The red dashed line indicates the earliest significant decrease in each region. (**d**) BOLD maps of the mPFC and thalamus at separate time points around LOC. (**e**) Cross correlations between the BOLD time series of thalamic ROIs and mPFC ROIs. Blue colors (i.e. negative time lags) indicate a leading phase from ROIs in the row to ROIs in the column. (**f**) BOLD intensity change around LOC in the hippocampus. (**g**) BOLD intensity changes around LOC in the brainstem.

To determine the exact time of deactivation, we conducted t-tests comparing the mean BOLD intensity in the thalamus and mPFC within 5-s bins from −50 s to 100 s to their corresponding baseline. Time bins with tests showing significantly reduced BOLD intensity compared to baseline (two-tail, p < 0.05) were marked in red in Figs. 3b,c,f. The earliest bin displaying significantly decreased BOLD intensity was identified and indicated by a red dashed line in Figs. 3b-c, f-g. As observed, the onset of deactivation in the thalamus precedes that of the mPFC by approximately 40-60 s. In addition, the recovery of the BOLD signal in the thalamus occurs earlier than in the mPFC (Fig. 3d, Fig. S5). Cross correlations between the BOLD time series of thalamic ROIs and mPFC ROIs confirm that the thalamic activity leads the mPFC activity (Figs. 3e).

Similarly to the thalamus and mPFC, the hippocampus and parahippocampus also exhibit a transient but smaller deactivation prior to LOC, with the onset time close to that of the thalamus and an earlier recovery time than both the thalamus and mPFC. However, the BOLD intensity of both the hippocampus and parahippocampus only recovers to approximately the baseline level eventually (see Fig. 3f and Fig. S6). This pattern was also observed in the averaged BOLD signal of the brainstem (Fig. 3g), although individual brainstem ROIs display relatively heterogeneous temporal profiles (Fig. S10), unlike the hippocampus, thalamus and mPFC.

In summary, these data suggest that there is a sequential process of neuronal activities, particularly along the thalamus-hippocampus-mPFC pathway, as the process of LOC unfolds. These results may indicate a traveling wave of brain activity surrounding LOC ^26,27,28,29^.

### Cortical regions exhibit elevated BOLD activity during sustained unconsciousness

While most cortical areas outside of the mPFC do not exhibit deactivation upon the onset of LOC, they do display a gradual, monotonic increase in BOLD signals thereafter (refer to Fig. 4a), each with varying ramp-up times before plateauing (see Fig. S7). To assess these changes statistically, we conducted t-tests comparing BOLD amplitudes between 300 s and 800 s after LOC (i.e., during sustained unconsciousness) to baseline levels. Overall, the cortex, excluding the mPFC, shows a significant elevation in BOLD amplitude during sustained unconsciousness (p = 0.009). Noteworthy cortical regions displaying suprathreshold BOLD amplitude during this period (Fig. 4d-e) include the primary motor cortex (M1, p = 0.0005), secondary motor area (M2, p = 0.017), secondary somatosensory area (S2, p = 0.047), entorhinal cortex (EC, p = 0.035), dysgranular insular cortex (DIC, p = 0.05) and granular insular cortex (GIC, p = 0.047). These findings suggest that alterations in cortical activity outside of the mPFC might be a consequence, rather than a cause, of LOC.

**Figure 4.**
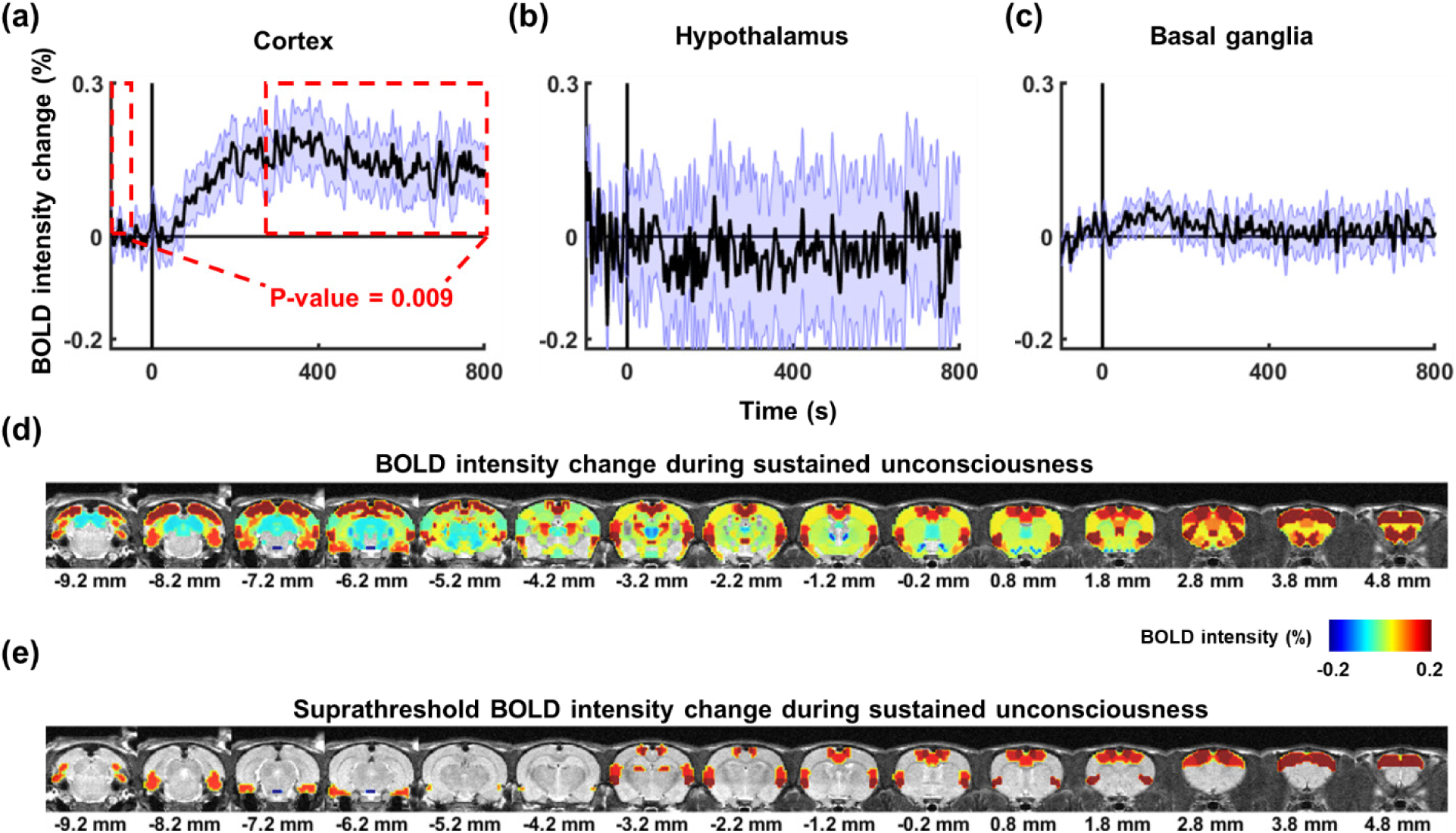
Brain-wide BOLD activity during sustained unconsciousness. (**a**) The cortex exhibits a gradual, monotonic increase in BOLD signal after the onset of LOC, reaching a plateau during sustained unconsciousness (defined as the period 300 s – 800 s post LOC, marked by the red dashed rectangle on the right), significantly exceeding the baseline level (the red dashed rectangle on the left, p = 0.009). (**b-c**) The **b**) hypothalamus and **c**) basal ganglia do not appear to be involved in the process of LOC. (**d**) Map of brain-wide BOLD intensity during sustained unconsciousness. (**e**) Suprathresholded map of (d) (p < 0.05).

Conversely, we did not observe appreciable changes in BOLD signals in the hypothalamus or basal ganglia, suggesting that these brain areas might not be involved in the process of LOC (Fig. 4b-e, Figs. S8, S9).

### Before the onset of LOC, there is a temporary increase in brain-wide functional connectivity

Given the similar temporal BOLD patterns observed across multiple brain regions such as the hippocampus, thalamus, and mPFC around the onset of LOC, we sought to investigate the alterations in brain-wide functional connectivity during LOC. To achieve this, we compared the dynamics of resting-state fMRI networks across three levels of consciousness: a baseline condition (representing the time before LOC, encompassing the initial 500 s under low-dose propofol), the time interval containing the onset of LOC (−100 s to 300 s around LOC), and the time period of sustained unconsciousness (300 s to 500 s after LOC).

BOLD time courses of individual ROIs were segmented using a 40-s sliding window with a 3-s step. For each window, the functional connectivity between every pair of ROIs was determined by calculating the Pearson correlation coefficient of their respective time courses. The resulting functional connectivity matrices from all windows across all rats were pooled, and subjected to k-means clustering into two clusters based on their brain-wide functional connectivity patterns. Additional clustering results for different cluster numbers (cluster number = 3 - 5) are illustrated in Figs. S11, 12, 13.

We evaluated the distribution of the occurrence rate of the two clustered patterns during both the baseline period and around the onset of LOC. For each pattern, the temporal distribution of each time window was quantified by calculating the ratio of occurrences of that pattern within a specific time window across all scans. The 95% confidence intervals, following a Bernoulli distribution, were determined using formula (1):

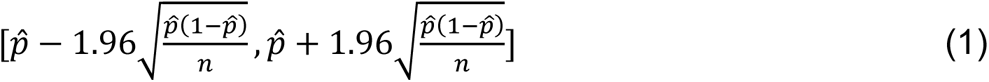

where 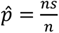 is the proportion of scans clustered into a given pattern; ns is the number of scans assigned to this pattern for a specific time window, and n is the number of total scans for this window.

To assess whether the temporal distribution of each pattern significantly changes around the onset of LOC compared to the baseline, two-tailed t-tests were conducted between the occurrence rates averaged across 15-s time bins segmented from −100 s to 100 s around LOC and under the low dose of propofol (i.e. baseline). Both patterns exhibited consistent temporal distributions under the baseline and the sustained unconsciousness state (300 s to 500 s after LOC) (two-tailed t-test, p-value = 0.6). However, relative to baseline, a higher occurrence was observed for pattern 2 from 50 s to 35s before LOC (two tailed t-tests, p_(−45s to −30s)_ = 0.041, uncorrected), and it showed a transient decrease from 55s to 65s after LOC (two tailed t-test, p(_45s to 60s_) = 0.027, uncorrected) (Fig. 5b). Pattern 1 showed the inverse change prior to LOC (see Fig. 5a), as the combined occurrence rate of the two patterns must equal 1.

**Figure 5.**
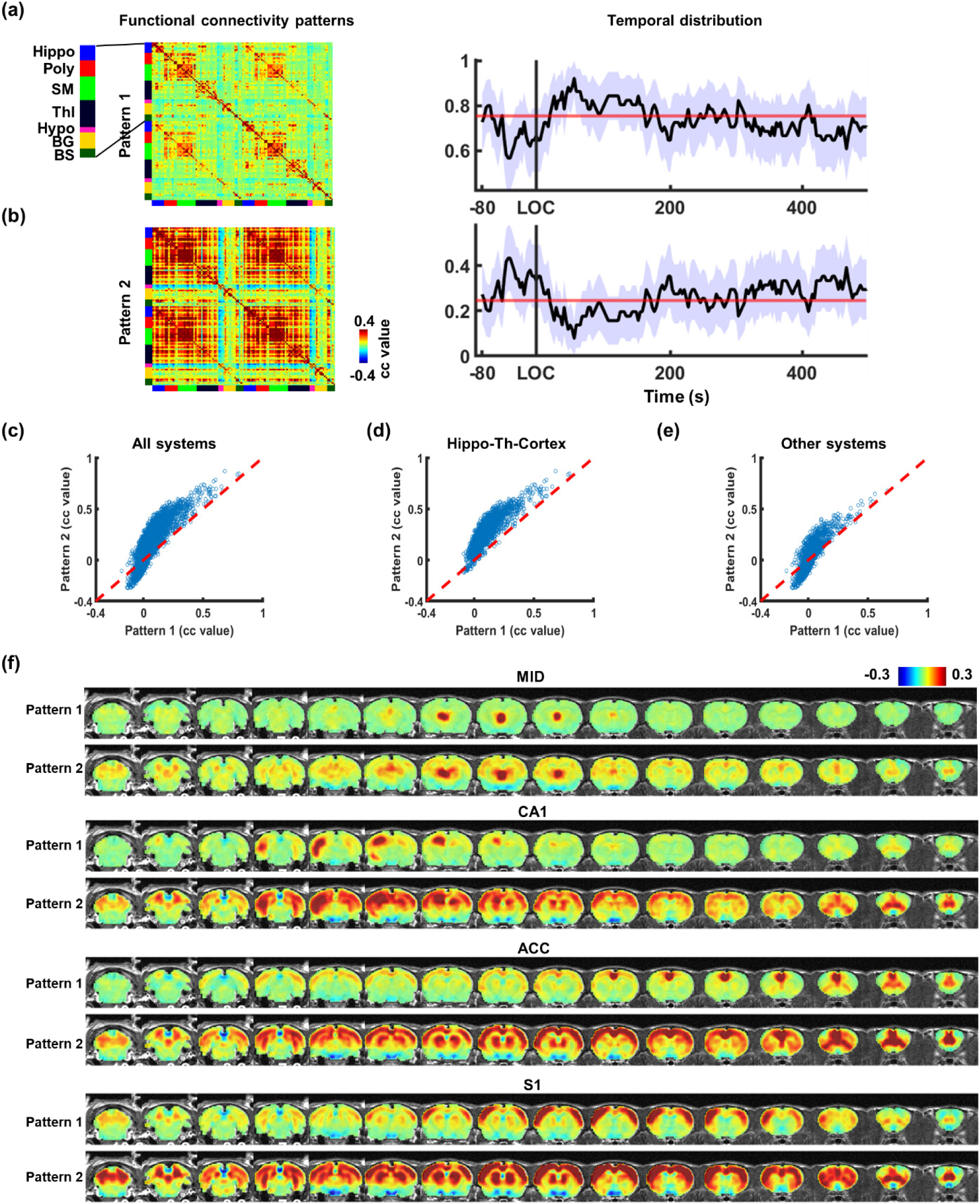
Dynamic brain states under different consciousness conditions. (**a,b**) Two brain states identified by unsupervised clustering of dynamic functional connectivity matrices, with their temporal distributions around the onset of LOC and during sustained unconsciousness. The purple shade indicates the 95% confidence interval. The red line represents the averaged temporal distribution under the low dose of propofol (i.e. baseline). Hippo: hippocampus; Poly: polynomial association cortex; SM: sensorimotor cortex; Thl: thalamus; Hypo: hypothalamus; BG: basal ganglia; BS: brainstem. (**c**) Comparison of functional connectivity strength between pattern 1 and pattern 2, with the red dashed diagonal line indicating where strengths are equal between the two patterns. (**d**) Comparison of functional connectivity strength in two patterns within the hippocampus-thalamus-cortex network. (**e**) Comparison of functional connectivity strength in two patterns in systems outside of the hippocampus-thalamus-cortex networks. (**f**) Averaged seed maps with the seeds located in the thalamus (MID), hippocampus (CA1), mPFC (ACC), and other cortical regions (S1) during time windows corresponding to the occurrence of either pattern.

Functional connectivity across the entire brain is distinct between pattern 1 and pattern 2. These differences were visualized in a scatter plot, with the x-axis depicting the functional connectivity strength in Pattern 1 and the y-axis indicating the functional connectivity of the corresponding connections in pattern 2. Overall, pattern 1 shows lower functional connectivity strength across the brain compared to pattern 2 (see Figs. 5c). Importantly, the heightened functional connectivity in pattern 2 is not uniform across the brain. Nearly all connections (99.6%) in the hippocampus-thalamus-cortex network display higher functional connectivity strength in pattern 2 compared to pattern 1 (see Fig. 5d), appreciably higher than other regions (Fig. 5e). Furthermore, for both patterns, seed maps of four ROIs in the left hemisphere, including the midline thalamic nucleus (MID, part of the thalamus), CA1 (part of the hippocampus), ACC (part of the mPFC), and primary somatosensory area (S1, part of other cortical regions), were generated for time windows when either pattern 1 or pattern 2 occurred, respectively. As depicted in Fig. 5f, all seed maps during the occurrence of pattern 2 showed higher and more widely distributed functional connectivity within the hippocampus-thalamus-cortex network compared to those during the occurrence of pattern 1. Taken together, in alignment with the transient BOLD intensity changes observed in the hippocampus-thalamus-mPFC pathway before LOC, our data also reveal increased functional connectivity within the same system during the same time window.

## Discussion

Utilizing simultaneous electrophysiology-rsfMRI recording, we investigated the spatiotemporal dynamics of brain activity during LOC. We first found that the timing of behaviorally determined LOC well aligns with the onset of low-frequency LFP power increase, consistent with previous literature indicating a rapid surge in slow fluctuations in cortical electrophysiology immediately following LOC^10,24,25,11,13^. Leveraging this electrophysiology signature of LOC, we identified a consistent temporal pattern of transit deactivation unfolding sequentially across the thalamus, mPFC, and hippocampus surrounding LOC. Conversely, BOLD activity in other cortical regions displayed gradual increases post LOC, eventually plateauing during sustained unconsciousness, suggesting that alterations in cortical activity outside of the mPFC might be a consequence, rather than a cause, of LOC. In addition, we noted a rise in BOLD synchronization along the hippocampus, thalamus, and cortex right preceding the LOC. These findings unveil a sequential cascade of brain activity that emerges as the animal transitions from consciousness to the unconsciousness, offering critical insight into the systems-level neural mechanisms underlying LOC.

The utilization of simultaneous electrophysiology-fMRI recording plays a key role in our current study. While it is widely recognized that LOC is characterized by abrupt behavioral changes and various neurophysiologic alterations, global changes in brain activity/connectivity during this rapid process remain elusive. To elucidate these changes, the simultaneous electrophysiology-fMRI method offers excellent capability. Electrophysiology recording has high temporal resolution, enabling precise determination of the timing of LOC, while fMRI provides high spatial resolution with a whole-brain field of view, facilitating the map of brain-wide activity/connectivity changes synchronized with LOC. Despite its technical challenges, our lab has successfully employed this method in our recent studies^17,18^. This capacity empowers us to comprehensively explore brain-wide dynamics during LOC.

A key discovery of our study unveils a cascade of transient deactivation occurring along the hippocampus-thalamus-mPFC pathway around the moment of LOC. Previous studies have consistently documented the association between deactivation of these three regions and LOC. For instance, Wang et al. reported that the paraventricular thalamic nuclei (PVT), a midline thalamic nucleus, begin to deactivate approximately 100 s before propofol-induced LOC^30^, closely aligning with our findings. In addition, chemogenetic or optogenetic stimulation of PVT neurons has been shown to reduce sensitivity to propofol and delay the onset time of LOC^30^. Similarly, reduced hippocampal activity has been observed under various anesthetics, including isoflurane, medetomidine-midazolam-fentanyl (MMF), and ketamine/xylazine-induced unconsciousness^31^. Chemogenetic deactivation of the hippocampus can lower the anesthetic dose required for inducing LOC and prolong the duration of unresponsiveness to stimuli during the recovery of consciousness^32^. Furthermore, deactivation of the prefrontal cortex accelerates the induction time of LOC^33^.

In line with these investigations, our data not only reaffirm the pivotal role of the hippocampus, thalamus, and mPFC in LOC, but also highlight a temporal lagging pattern among these regions during LOC, implying that the interaction in the hippocampus-thalamus-mPFC network might be key to triggering LOC. This notion is further supported by an increased occurrence of synchronization in their BOLD activity before LOC and a subsequent decrease after LOC in our study, as well as in several previous studies. For example, research has shown that the activation of the mPFC by phencyclidine, a hallucinogen that can alter the state of consciousness, is mediated via the hippocampo-prefrontal pathway^34^. The dynamic hippocampal-mPFC connectivity observed in awake animals is diminished in a steady anesthetized state^35^. Additionally, the level of consciousness can be altered by thalamic stimulation, accompanied by changes in coherence between the deep layer of the prefrontal cortex and the superficial layer of the parietal cortex^36^. Considering that the deep cortical layers send feedback to superficial layers in lower-order cortical areas^37^, this result may suggest that the thalamus can modulate consciousness levels by altering signal propagation from prefrontal cortex to lower-order cortical regions. Taken together, alterations in coordinated activity within the hippocampus-thalamus-mPFC pathway are tightly linked to LOC. Based on these findings, we conjecture that brain-wide activity propagation may be disrupted by the sequential deactivation of the hippocampus, thalamus and mPFC, leading to LOC. This speculation agrees with the concept that hyper-synchrony across the brain could disrupt effective communication, and thus trigger the transition from consciousness to unconsciousness^38^.

Besides the cascade of transit deactivation in the hippocampus, thalamus and mPFC around the onset of LOC, our study showed a gradual increase in the BOLD signal in multiple cortical regions, eventually reaching an above-baseline plateau post-LOC. This result well agrees with the strong vasodilation observed during sleep reported in animal studies using fMRI and optical imaging^39,40^. This change could be triggered by the diminished release of vasoconstrictive norepinephrine (NE) from the locus coeruleus during AIU^41^ and sleep^42,43^, given its widespread cortical projections. Furthermore, the increased synchronization in the cortex during AIU, as indicated by calcium imaging data^44^, may also contribute to the increase of the BOLD signal^45^. These results together suggest that heightened blood flow/blood volume/BOLD activities in the cortex might constitute a general phenomenon across various forms of steady-state unconsciousness. Importantly, all these changes occurred tens to hundreds of seconds after LOC during sustained unconsciousness, suggesting that the BOLD increase observed in these cortical regions might be a resultant effect, rather than a cause of LOC.

It is likely that the observed change in global synchrony, as revealed by fMRI data, results from variations in motion and/or other physiological factors surrounding LOC^46^. To rule out the possibility, in rsfMRI data preprocessing we regressed out motion parameters and the signals of the white matter and cerebrospinal fluid (CSF)^47^. In addition, we observed that both respiration, quantified by respiration volume per time (RVT), and motion level remained stable prior to LOC, and both decreased post-LOC (Figs. S14-S15). This stability suggests that changes detected in electrophysiology/rsfMRI signals around the onset of LOC are not predominantly influenced by these non-neural factors.

## Conclusion

We identified a sequence of brain activity unfolding across the hippocampus, thalamus, and mPFC during transitions from consciousness to unconsciousness. Our findings not only reaffirmed the important role of these three systems in consciousness state transitions but also uncovered a temporal pattern intricately linked to the moment of LOC. Moreover, we observed transient fluctuations in long-distance synchronization across the brain during LOC. Overall, these results underscore the pivotal role of coordinating hippocampus, thalamus, and mPFC activities both spatially and temporally during LOC. This study also highlights the value of simultaneous electrophysiology-rsfMRI recording in elucidating the changes of whole-brain activity during state transitions.

## Methods

### Animals

Sixteen adult male Long-Evans rats (400 - 650 g) were used in this study. Nine rats were included in imaging experiments and seven rats were involved in behavior experiments. Animals were housed in Plexiglas cages with ad libitum access to water and food. The room temperature was maintained at 22–24^°C^ under a 12-hour light: 12-hour dark cycle. All experiments were approved by the Pennsylvania State University Institutional Animal Care and Use Committee (IACUC).

### Surgery

Nine animals underwent aseptic surgeries for the chronic implantation of electrodes. Anesthesia was initiated with isoflurane, followed by the intramuscular administration of a ketamine-xylazine cocktail (40 mg/kg and 12 mg/kg respectively). Baytril (2.5 mg/kg) was administered subcutaneously as an antibiotic. Postoperative analgesia was provided through subcutaneous injection of Buprenorphine (1.0 mg/kg). Throughout the surgery, the body temperature was maintained at 37^°C^ using a warming pad (PhysioSuite, Kent Scientific Corporation). Vital parameters, including oxygen saturation (SpO_2_) and heart rate, were continuously monitored with a pulse oximetry (MouseSTAT Jr, Kent Scientific Corporation). During the surgery, a constant supply of a gas mixture of oxygen and isoflurane (0–2%) was provided through a nose cone to maintain the anesthetic state, and the animals were immobilized on a stereotaxic platform (David Kopf Instruments, Tujunga, CA). A craniotomy was made over the left anterior cingulate cortex (ACC, coordinates: anterior/posterior +1.5, medial/lateral −0.5, dorsal/ventral −2.2), and an MR-compatible 16-channel electrode (MRCM16LP, NeuroNexus Inc) was carefully inserted into the ROI. The electrode’s reference and grounding wires were securely attached to a silver wire positioned on the surface of the right cerebellum. The electrode was then affixed to the skull using a tissue adhesive (Vetabond, 3M, St. Paul, MN) and dental cement (ParaBond, COLTENE, Cuyahoga Falls, OH). After surgery, rats were returned to a clean home cage, and post-surgery care was provided for at least 7 days. During this recovery period, animals were not subjected to any experimental procedures.

### Simultaneous rsfMRI and electrophysiology recordings

rsfMRI was performed on a 7T Bruker 70/30 BioSpec scanner through ParaVision 6.0.1 software (Bruker, Billerica, MA) at the *high field MRI facility* at the Pennsylvania State University. rsfMRI data were collected using a T_2_*-weighted gradient-echo echo-planar-imaging (EPI) sequence with the following parameters: echo time (TE) = 15 ms; repetition time (TR) = 1000 ms; in-plane resolution = 0.5 × 0.5 mm^2^; slice thickness = 1 mm; number of slices = 20; FOV = 32 × 32 mm^2^; image matrix = 64 × 64.

Prior to imaging, the rats were set up in a custom-designed rodent restrainer^48^, with artificial tears applied to their eyes to prevent dryness, and ear plugs to protect the rats from the scanning noise. Before rsfMRI scanning, a bolus injection of propofol (20 mg/kg) was administered to the rat through intravenous injection, then a low dose of propofol (20 mg/kg/h) was infused for at least 40 minutes to achieve a steady state before the first rsfMRI scan (10 mins long) was acquired. Subsequently, the propofol dose was increased to 40 mg/kg/h for 20 minutes, as the second rsfMRI scan was continuously collected during this period. The paradigm is schematically illustrated in (Fig. 1a). Throughout all imaging sessions, the animal’s core temperature was continuously monitored by a rectal thermometer and maintained at 36.5 – 37.5^°C^ through a water warming system (GAYMAR T/ PUMP TP420). The rodent’s respiration signal was recorded via a respiration sensor placed under the animal’s chest with the sampling rate of 225 Hz. Throughout the rsfMRI scanning process, the electrophysiology signal was simultaneously recorded from the left ACC using a TDT recording system (Tucker Davis Technologies (TDT) Inc, Alachua, FL). The raw electrophysiology signal was collected via an MR-compatible LP16CH headstage, then amplified and digitized in a PZ5 neurodigitizer amplier (TDT Inc, Alachua, FL) and recorded using Synapse software and an RZ2 BioAmp Processor (TDT Inc, Alachua, FL). The raw electrophysiology signal was sampled at either 24414 Hz (45 scans in total) or 6104 Hz (6 scans in total) and stored on a WS8 workstation (TDT Inc, Alachua, FL).

### Behavioral assessment of unconsciousness

The consciousness level of the rat in relation to the dose and timing of propofol administration was tested using the loss of righting reflex (LORR) test, a well-established method for determining the level of consciousness^49^. This test was conducted outside the scanner in 7 rats, following the same anesthesia protocol as used during scanning and accompanied by prerecorded EPI scanning noise to mimic the imaging environment. The LORR test was conducted by rapidly placing the rat in a supine position, and a complete LORR was determined if the rat failed to return to the prone position within 30 s. Each LORR experiment is comprised of 4 trials: the first trial occurs under the steady state of low-dose propofol, at 5 min before the onset of high dose; the 2^nd^, 3^rd^ and 4^th^ trials occurred at the 2-min, 5-min and 10-min marks following the start of high-dose administration, respectively.

### Data preprocessing

An in-house, MATLAB-based preprocessing pipeline was used for rsfMRI data preprocessing, which is described in detail elsewhere^50^. Briefly, rsfMRI images were first motion scrubbed based on framewise displacement (FD) values. To reduce motion artifacts, volumes with FD > 0.1 mm were excluded along with their immediately adjacent volumes. To ensure MR signal stability, the first 10 volumes of each scan were discarded. Scans with more than 10% of volumes removed were not included in further analysis. After motion scrubbing, EPI images were co-registered to a predefined structural image template, and then underwent motion correction, nuisance regression of six motion parameters and the averaged BOLD signal from voxels in the white matter and CSF, independent component analysis cleaning, spatial smoothing (FWHM = 1 mm), and temporal filtering (0.001-0.1 Hz) with the mean BOLD intensity of each voxel added back after regression.

The electrophysiology signal recorded during rsfMRI scanning is comprised of a signal mixture derived from neural activity, electrode-specific noise, and MR-induced artifacts^51^. To remove MR-induced artifacts and denoise raw electrophysiology data, the following preprocessing pipeline was implemented. Firstly, the electrophysiology signal was segmented and aligned to the corresponding rsfMRI scans. Then phase correction across all 16 channels was conducted by cross correlation. Phase-corrected signals from all 16 channels were averaged together to create a single time series for each scan. This time series was then used to segment the scan into intervals corresponding to each individual fMRI slice acquisition. The time stamps of all individual segments were then applied to the electrophysiology signal of individual channels. For each channel, an MRI artifact template specific to each fMRI slice (i.e. segment) was determined using the average of the channel’s electrophysiology data across the first 400 segments (i.e. 20-second window). The MR-induced artifacts were then removed from raw electrophysiology signal by linearly regressing out the artifact template. This process was repeated for all segments across all channels. Following this, the resulting electrophysiology signal underwent notch filtering to remove power supply noise (60 Hz and its harmonics) and slice acquisition noise (20 Hz and its harmonics). After that, PCA denoising was further applied to the signal in the high-frequency band (> 150 Hz). Finally, the LFP was extracted using a bandpass filter (0.2 – 100 Hz). The quality of denoised LFP was shown in Fig. S2. Cross correlations between the BOLD signal of the left ACC and HRF-convolved (HRF: a single gamma probability distribution function [a = 3, b = 0.8]^52^) LFP powers were calculated with time lags ranging from −20s to 20s (Fig. 1b).

### Data Analysis: electrophysiology

The power spectrum of denoised electrophysiology signal was calculated using a multi-taper spectral estimation (taper number = 3, window size = 3 s, step size = 0.5 s) supplied by the Chronux toolbox (http://www.chronux.org). We calculated band-limited powers for different frequency ranges: slow wave (0.2-1 Hz), delta (1–4 Hz), theta (5–7 Hz), alpha (8–12 Hz), beta (13–30 Hz), and gamma (40–100 Hz), by averaging the power of all frequency bins within each frequency band. Our preliminary analysis revealed a noticeable upwards shift in the low-frequency band power after the initiation of administration of high-dose of propofol (Figs. 1d-e). Therefore, we manually identified the onset of low-frequency LFP power increase based on the powers in the slow wave (0.2-1 Hz) and delta (1-4 Hz) band. The power change temporally proximal to this onset (from 100 s before onset to 300 s after onset) was normalized as the percentage change relative to the baseline period from 100 s to 10 s before the onset. To examine how band-limited powers changed after the manually selected onset, for each frequency band, the LFP power of each 10-second time window, starting from 20 s before the onset to 800 s after the onset with no overlap between windows, was compared to the LFP power of baseline, defined as the time window from 120 s to 10 s before the onset to the onset. These comparisons used two sample, two-tailed T-tests, with p-value < 0.05 used as the statistical significance threshold.

### Associate the onset of low-frequency LFP power increase with LORR

After identifying the onset of low-frequency LFP power, we calculated the ratio of total scans whose low-frequency power increase took place at 0 min, within 2 min, within 5 min, and within 10 min, respectively, following high-dose propofol administration. Subsequently, Pearson correlation was conducted between this ratio and the ratio of animals exhibiting LORR in the corresponding time intervals. This test assessed the extent to which the onset of low-frequency LFP power increase aligns with the LORR in the same time window, and therefore helps determine whether the onset of low-frequency LFP power increase can be used as an electrophysiological proxy of LOC determined by the behavioral test.

### Data Analysis: rsfMRI

To investigate spatiotemporal dynamics of brain-wide activity around LOC, we specifically examined rsfMRI data segment from 100 s before to 800 s after the electrophysiology signature-informed LOC (900 sec in duration). To control for any potential influence of motion in this analysis, any segment having more than 20 rsfMRI volumes during the time intervals from −100 s to 100 s around LOC (i.e. 10% volumes) with FD > 0.1 mm were excluded from this analysis. As LOC always took place in the 2^nd^ EPI scan of each imaging session, during which the propofol dose was increased from 20 mg/kg/h to 40 mg/kg/h, for segments in scans with electrophysiology-informed LOC earlier than 100 s into the 2^nd^ scan, all missing time points were pre-padded with NAN (i.e. not-a-number). For statistical calculations, NAN values were treated as missing data, and regions with distortion were excluded, which were identified as areas with low BOLD intensity. To analyze activity changes in specific regions, the entire brain was parcellated into 59 bilateral regions of interest (ROIs) and categorized into 8 brain systems based on anatomical definitions provided in the Swanson Atlas^53^. For each scan, the ROI time courses were computed as the averaged BOLD time courses from all voxels within each ROI. The BOLD time series for these ROIs were then normalized as the percentage change relative to the same ROI’s average BOLD intensity during −100 s to −45 s relative to the LOC. To ensure the length of baseline is at least 5s, scans with onset earlier than 50s after the initiation of high dose propofol were excluded from further analysis. The resulting scan-specific normalized BOLD time series were time locked to the moment of LOC, and then averaged across all scans.

### Dynamic functional connectivity

Dynamic changes in brain functional connectivity surrounding LOC were analyzed using a sliding window approach (window length = 40 s, step size = 3 s). These specific parameters were chosen based on findings from previous studies, and alternative window length and step size also yielded similar results^54^. Due to a potentially variable distribution of dynamic functional connectivity (dFC) patterns, which might be dependent on steady consciousness levels or the transition between them, we analyzed the following sections of rsfMRI data: the first 500 seconds under low-dose propofol (45 scans in total) and the −100 to 500 s interval around the LOC under high-dose propofol (51 scans in total).

Restricting the sliding windows to the aforementioned intervals, 323 windows were generated and the dynamic functional connectivity matrix for each window was calculated as the Pearson correlation coefficient between the within-window time courses of each pair of 118 unilateral ROIs. At this point, for a given scan and a given window, a 118 x 118 symmetric correlation matrix was generated. This procedure was repeated for all windows and all scans. K-means clustering was then applied to this collection of dFC matrices. The K-means algorithm was applied using several cluster sizes: from 2 to 5. Ultimately, a cluster size of 2 was chosen because cluster sizes of 3-5 did not offer any additional useful information in extracting dFC patterns (Figs S11-13).

The RVT was determined as the difference between two adjacent peaks of the respiration signal divided by the duration between these two peaks.

## Acknowledgments

The present study was partially supported by the National Institute of General Medical Sciences (R01GM141792) The content is solely the responsibility of the authors and does not necessarily represent the official views of the National Institutes of Health.

