## Supplemental Document for "Sequential deactivation across the thalamus-hippocampus-mPFC pathway during loss of consciousness"

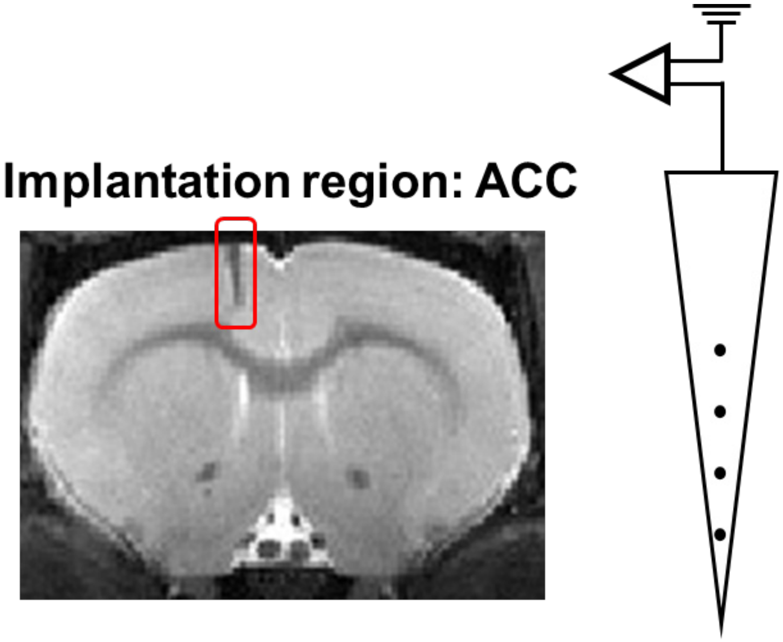


**Figure S1. An exemplar T2-weighted image showing an MR-compatible electrode implanted at rat's left ACC.**

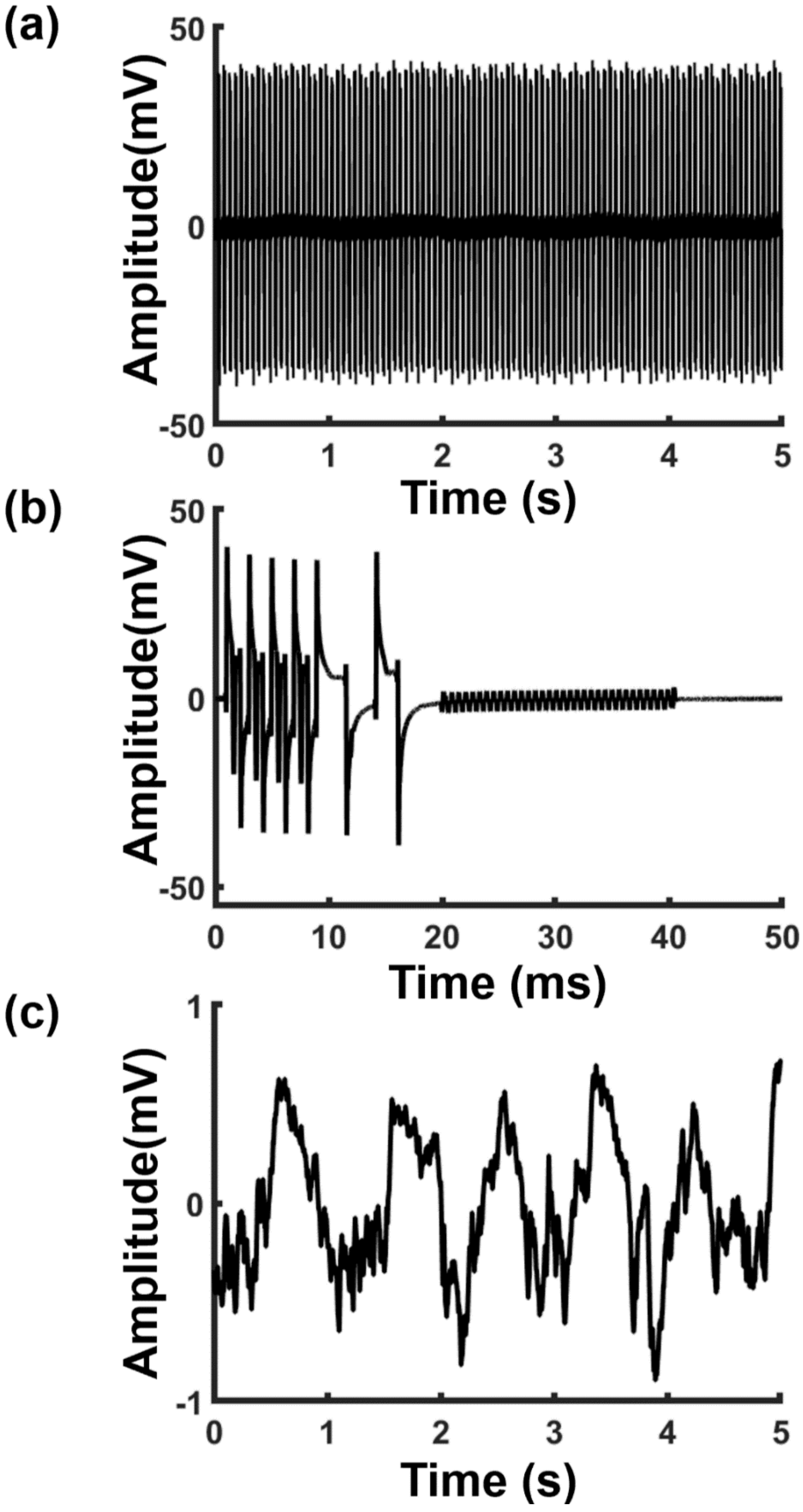


**Figure S2. Denoising MR-induced artifacts from electrophysiology signals.** (**a**) Raw electrophysiology data with MR-induced artifact in a 5-s interval. (**b**) A template of MR-induced artifact. (**c**) Denoised local field potential (LFP) data of the same 5-s interval as (a).

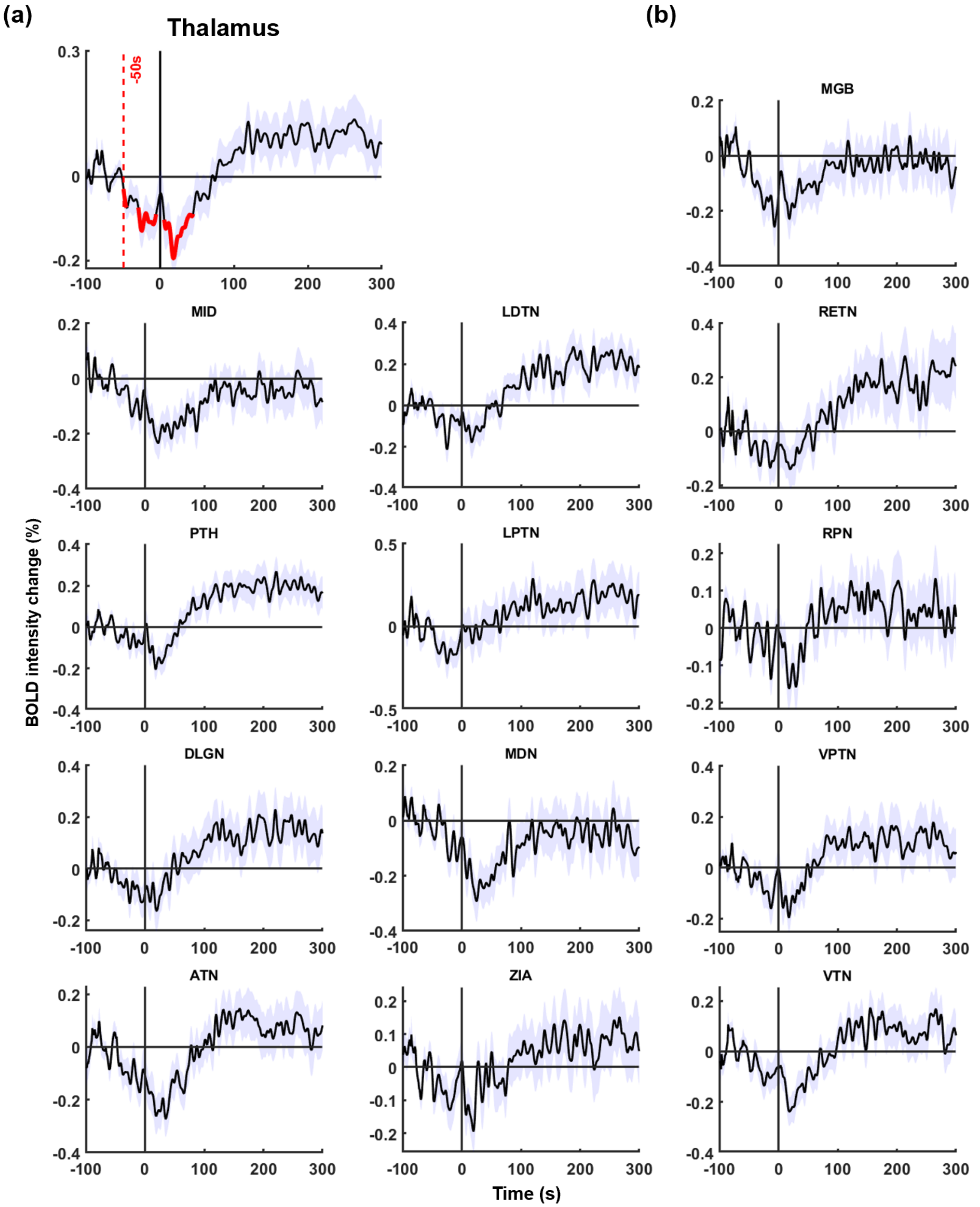


**Figure S3. BOLD intensity changes around the onset of LOC in subregions of the thalamus.** (**a**) BOLD intensity change in the thalamus (the same as Fig. 3b). (**b**) BOLD intensity changes of individual nuclei within the thalamus. From left to right, top to bottom: medial geniculate body (MGB), midline thalamic nucleus (MID), laterodorsal thalamic nucleus (LDTN), reuniens thalamic nucleus (RETN), posterior thalamic nucleus (PTH), lateral posterior thalamic nucleus (LPTN), reticular (pre)thalamic nucleus (RPN), dorsal lateral geniculate nucleus (DLGN), mediodorsal thalamic nucleus (MDN), ventral posterior thalamic nucleus (VPTN), anterior thalamic nucleus (ATN), zona incerta (ZIA), and ventral thalamic nucleus (VTN).

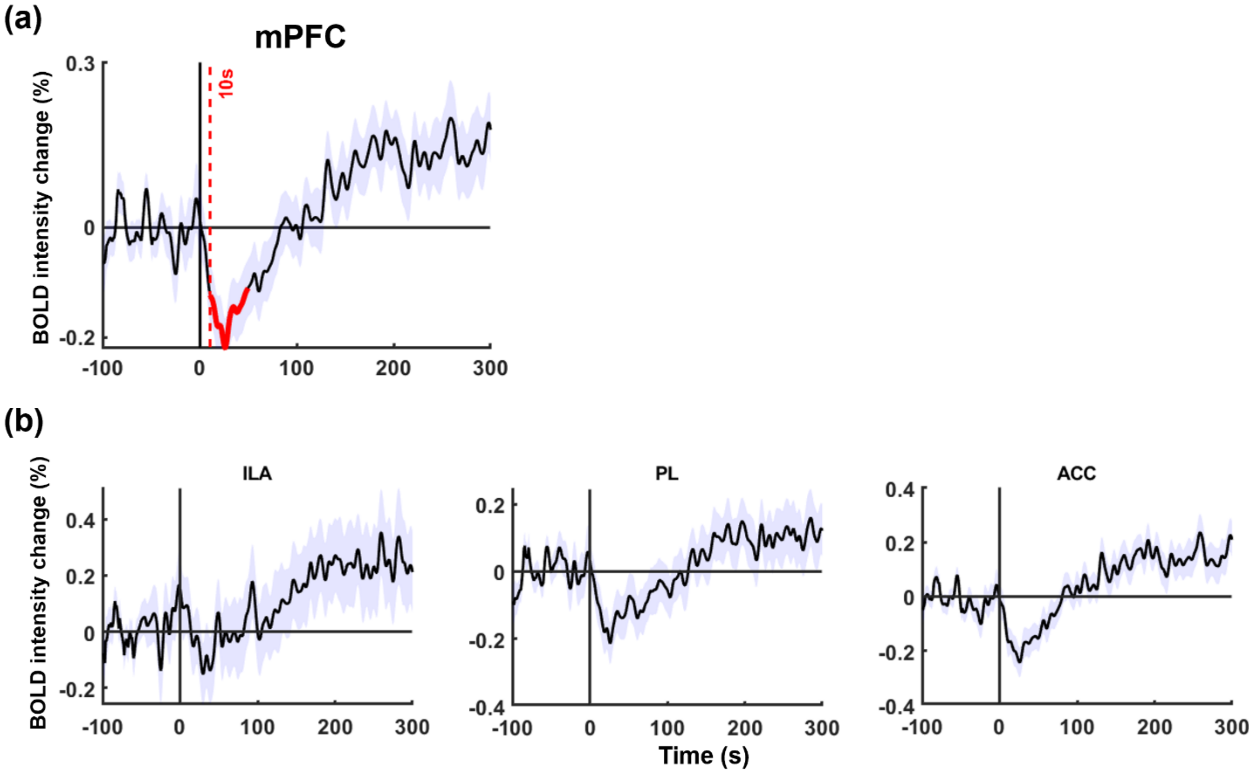


**Figure S4. BOLD intensity changes around the onset of LOC in subregions of the medial prefrontal cortex.** (**a**) BOLD intensity change in the mPFC (same as Fig. 3c). (**b**) BOLD intensity changes of individual subdivisions within the mPFC. From left to right: infralimbic area (ILA), prelimbic area (PL), and anterior cingulate cortex (ACC).

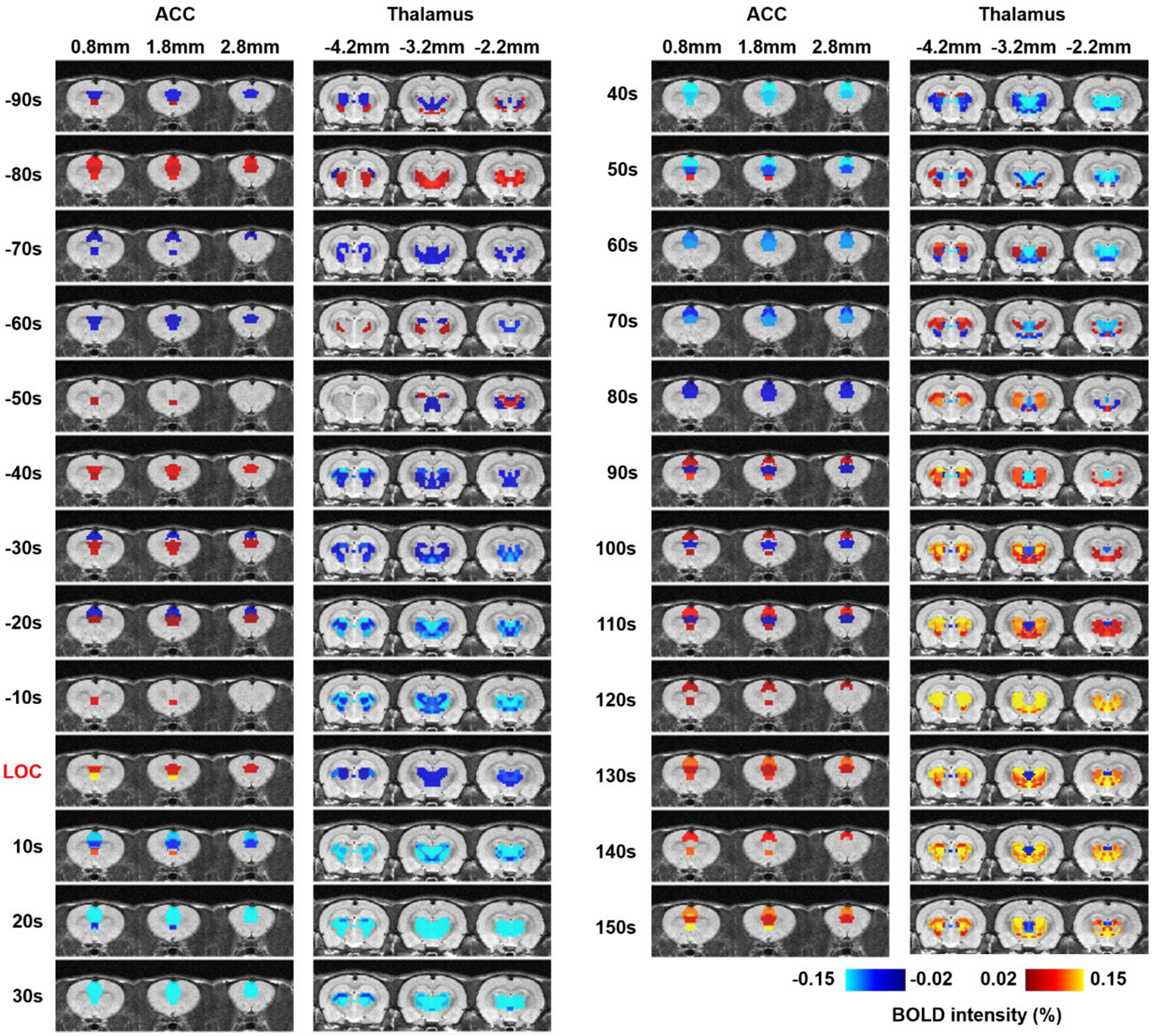


**Figure S5. BOLD intensity changes of ROIs in the thalamus and mPFC within the time window of -90s to 150s around the onset of LOC.** The BOLD intensity is normalized to the baseline level, defined as the averaged BOLD intensity during the period of -100s to -45s before LOC.

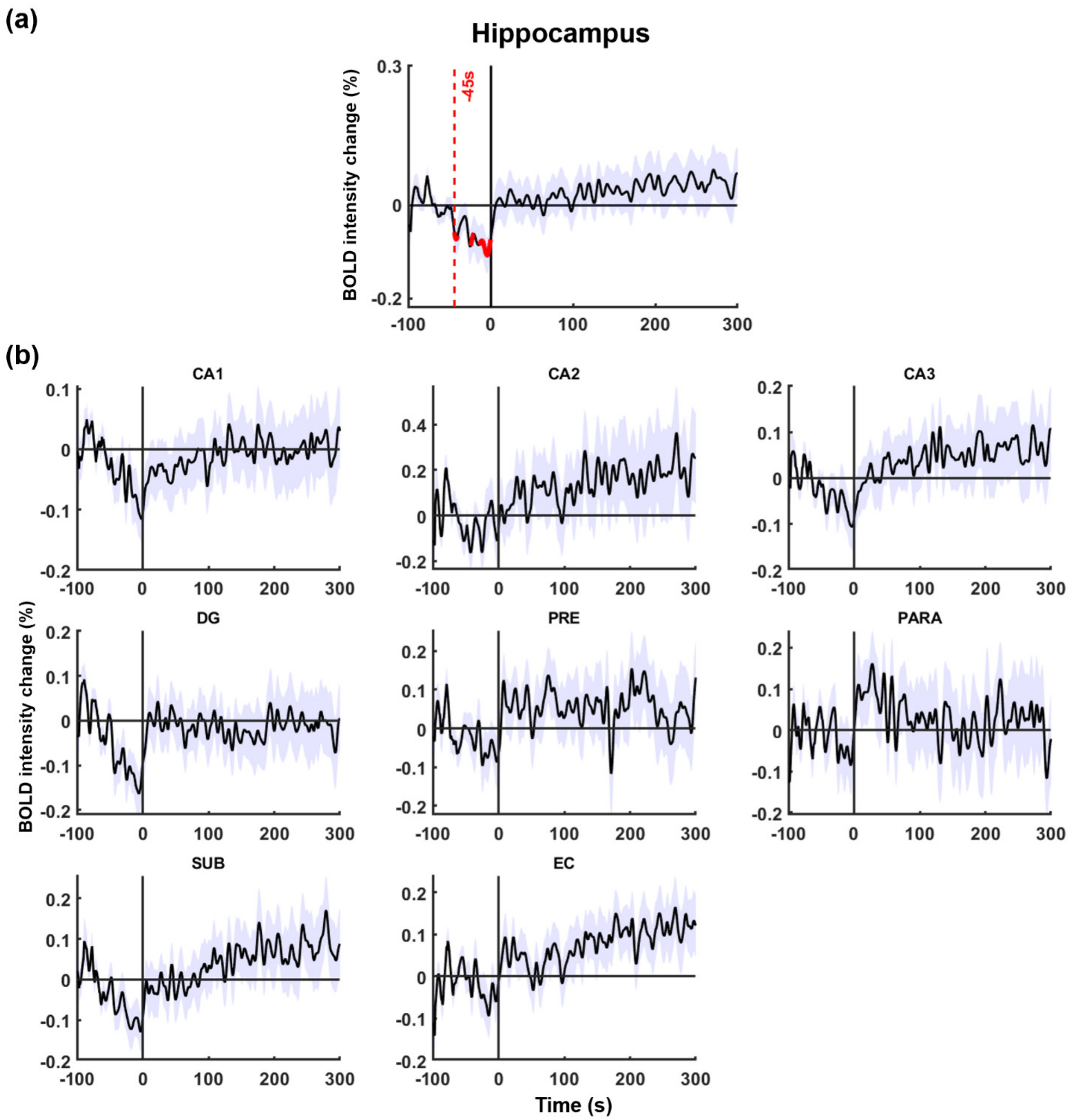


**Figure S6. BOLD intensity changes around the onset of LOC in subregions of the hippocampus.** (**a**) BOLD intensity change in the hippocampus (the same as Fig. 3f). (**b**) BOLD intensity changes of all subregions within the hippocampus. From left to right, top to bottom: cornu ammonis 1 (CA1), cornu ammonis 2 (CA2), cornu ammonis 3 (CA3), dentate gyrus (DG), presubiculum (PRE), parasubiculum (PARA), subiculum (SUB), and entorhinal cortex (EC).

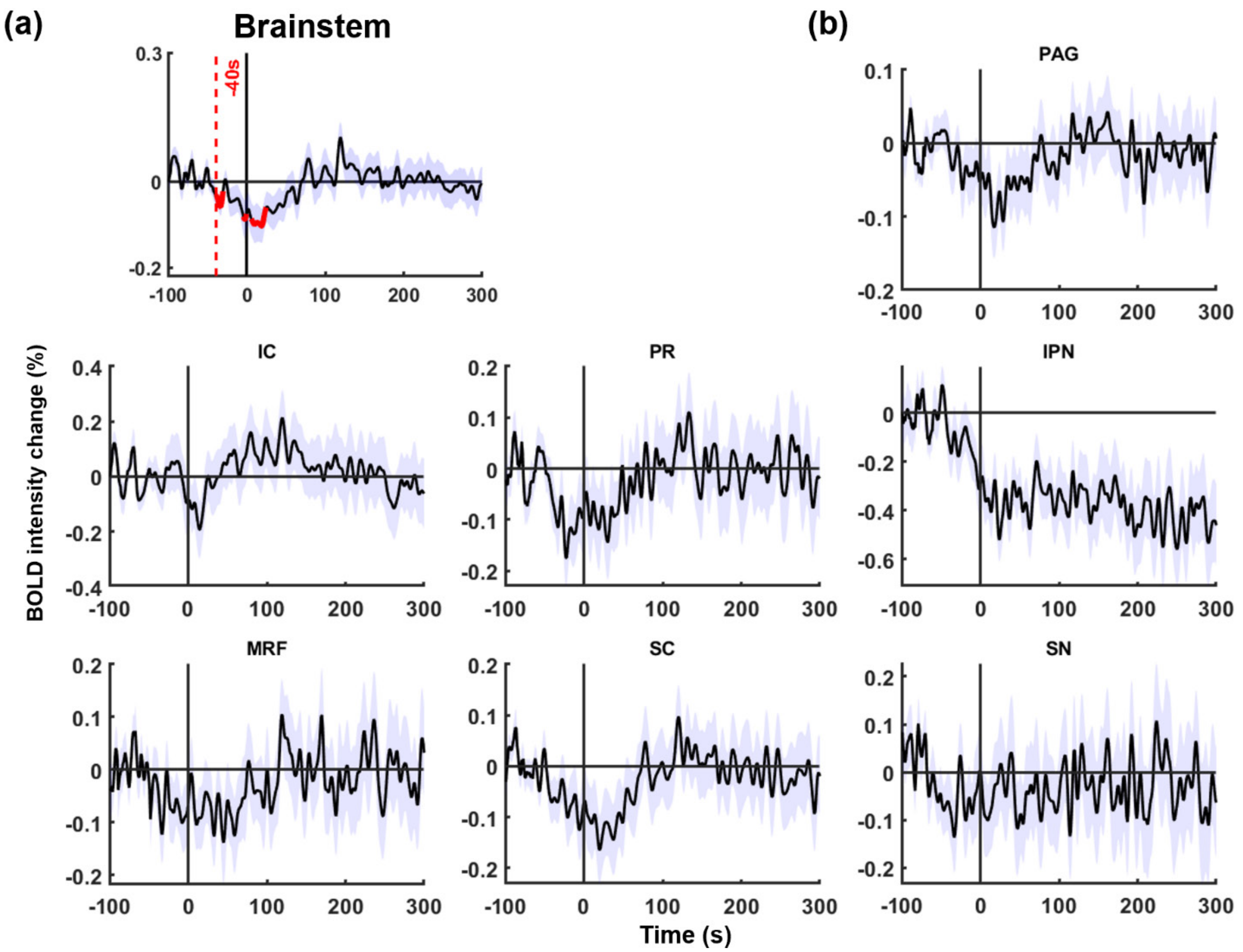


**Figure S7. BOLD intensity changes around the onset of LOC in subregions of brainstem.** (**a**) BOLD intensity change in the brainstem (the same as Fig. 3g). (**b**) BOLD intensity changes of individual subregions within the brainstem. From left to right, top to bottom: periaqueductal gray (PAG), inferior colliculus (IC), pretectal region (PR), interpeduncular nucleus (IPN), mesencephalic reticular formation (MRF), superior colliculus (SC), and substantia nigra (SN).

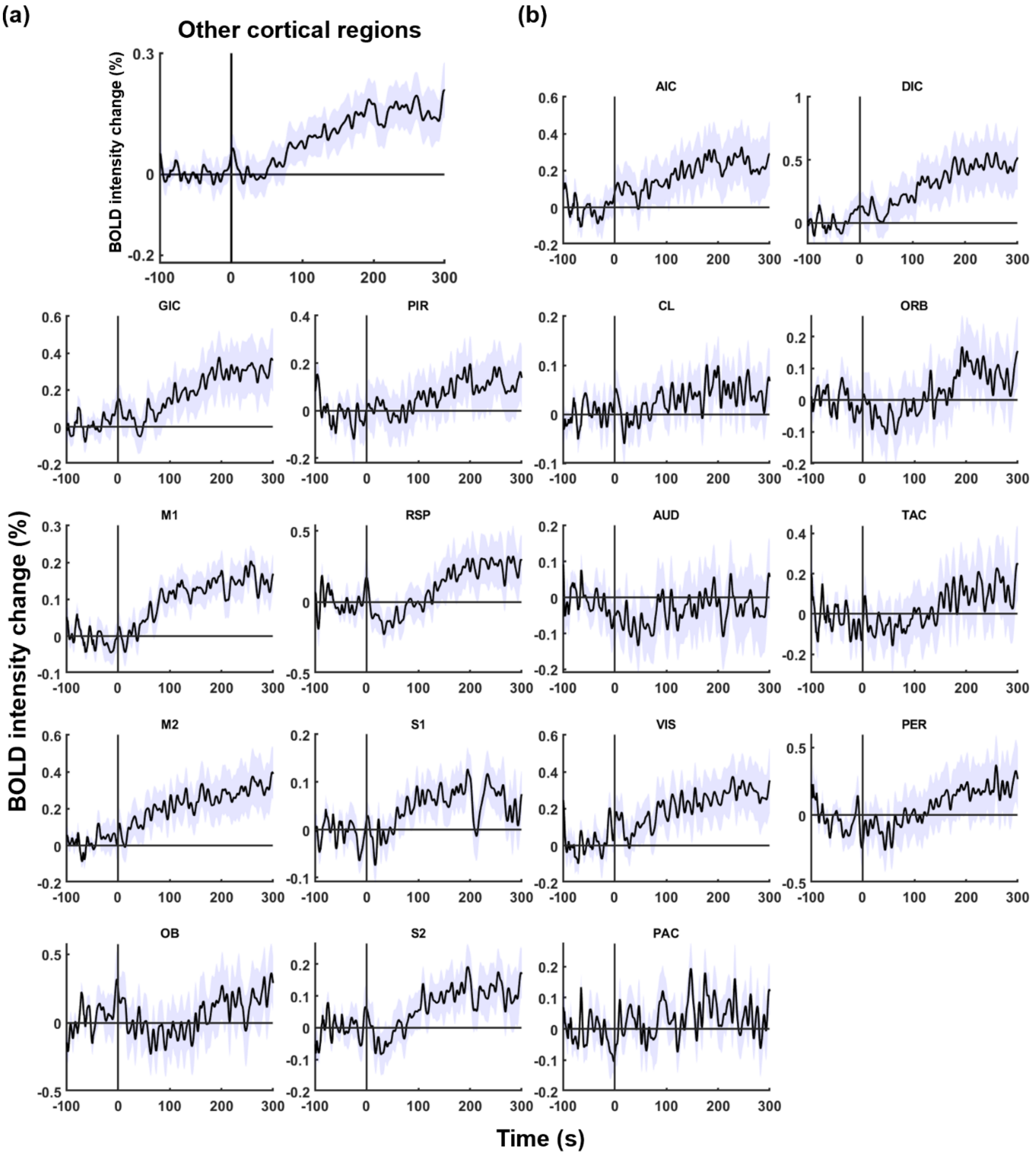


**Figure S8. BOLD intensity changes around LOC in other cortical regions.** (**a**) BOLD intensity change in cortical regions excluding the mPFC. (**b**) BOLD intensity changes of all ROIs included in the cortex except for the mPFC. From left to right, top to bottom: Agranular insular cortex (AIC), Dysgranular insular cortex (DIC), Granular insular cortex (GIC), Piriform cortex (PIR), Claustrum (CL), Orbital area (ORB), Primary motor area (M1), Retrosplenial area (RSP), Auditory area (AUD), Temporal association cortex (TAC), Secondary motor area (M2), Primary somatosensory area (S1), Visual area (VIS), Perirhinal area (PER), Olfactory bulb (OB), Secondary somatosensory area (S2), Parietal association cortex (PAC).

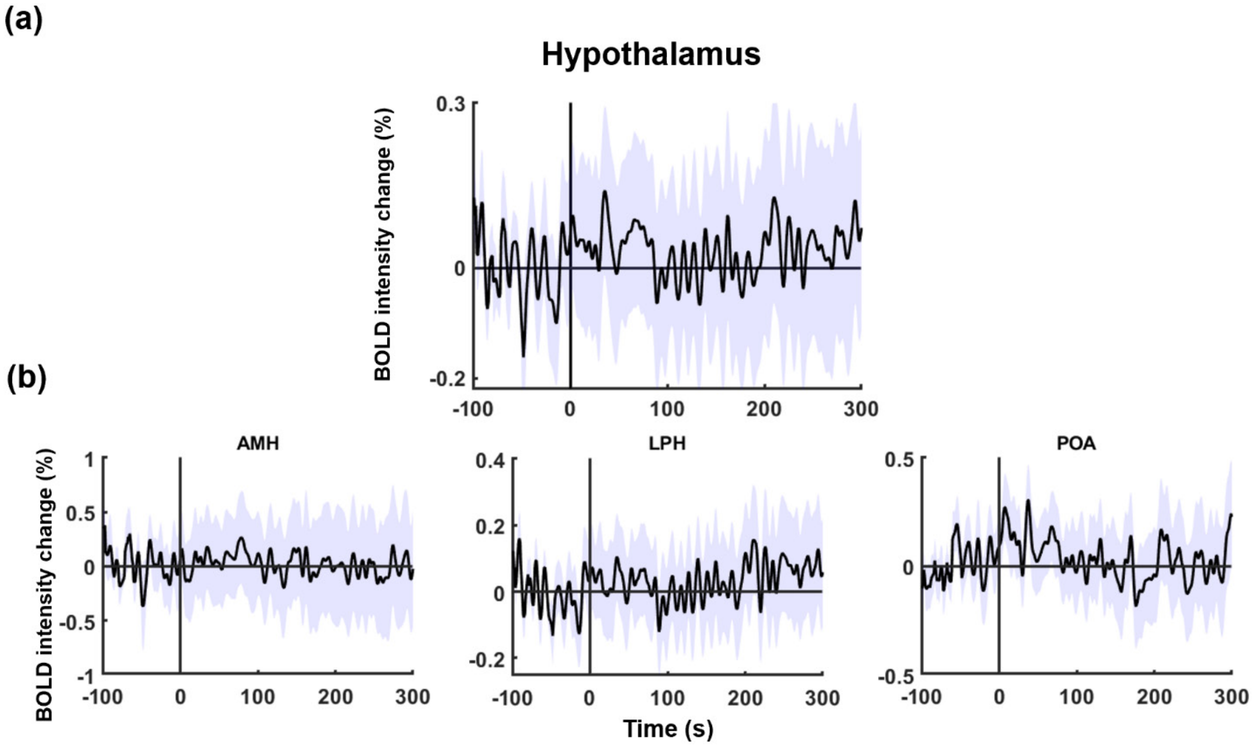


**Figure S9. BOLD intensity changes around LOC in subregions of hypothalamus.** (**a**) BOLD intensity change in the hypothalamus. (**b**) BOLD intensity changes of all ROIs included in hypothalamus. From left to right: Anterior medial hypothalamus (AMH), Lateral posterior hypothalamus (LPH), Preoptic area (POA).

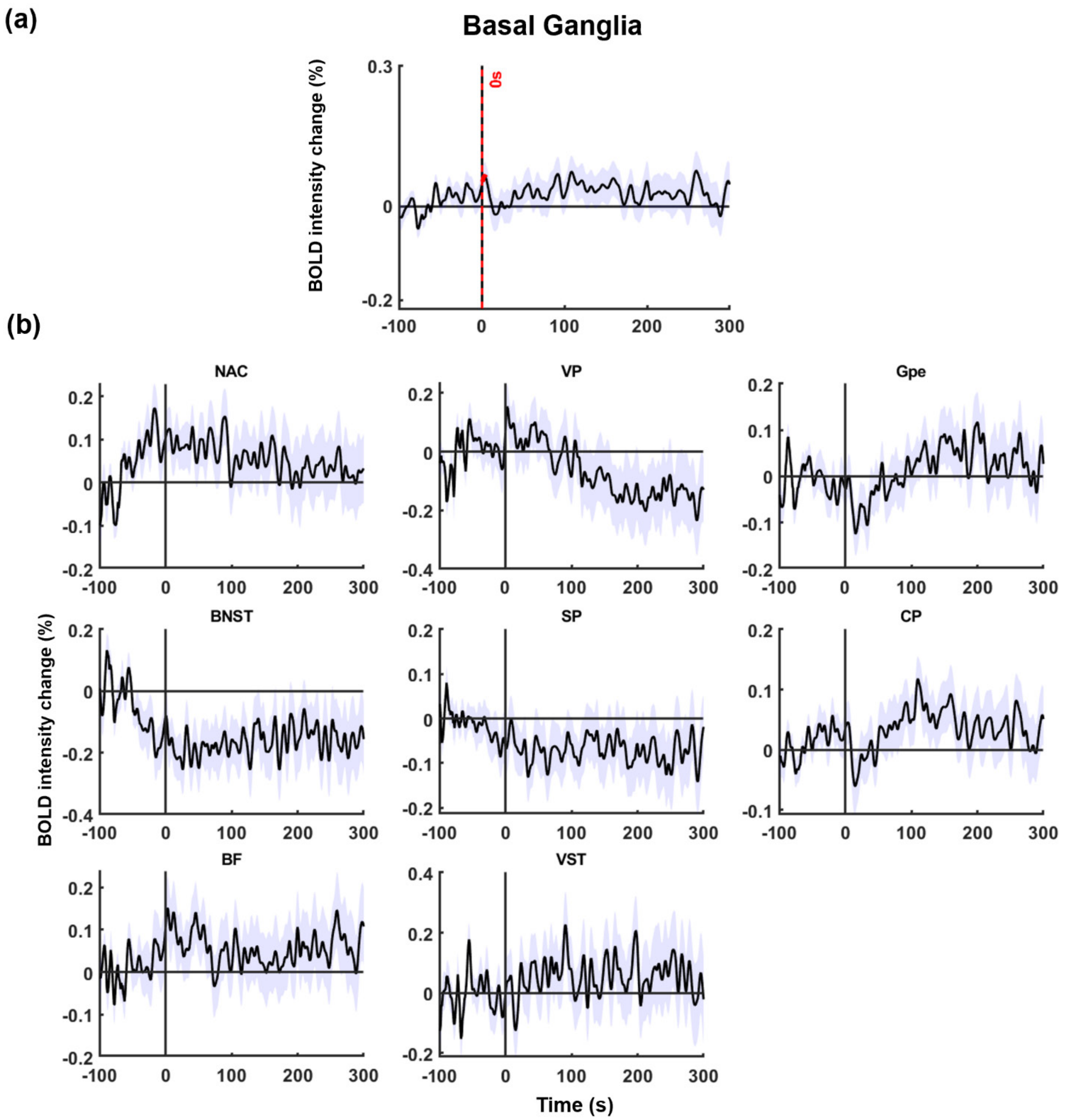


**Figure S10. BOLD intensity changes around LOC in subregions of basal ganglia.** (**a**) BOLD intensity change in the basal ganglia. (**b**) BOLD intensity changes of all ROIs included in basal ganglia. From left to right, top to bottom: Nucleus accumbens (NAC), Ventral pallidum (VP), Globus pallidus (Gpe), Bed nucleus of the stria terminalis (BNST), Septal region (SP), Caudate putamen (CP), Basal forebrain (BF), Ventral striatal region (VST).

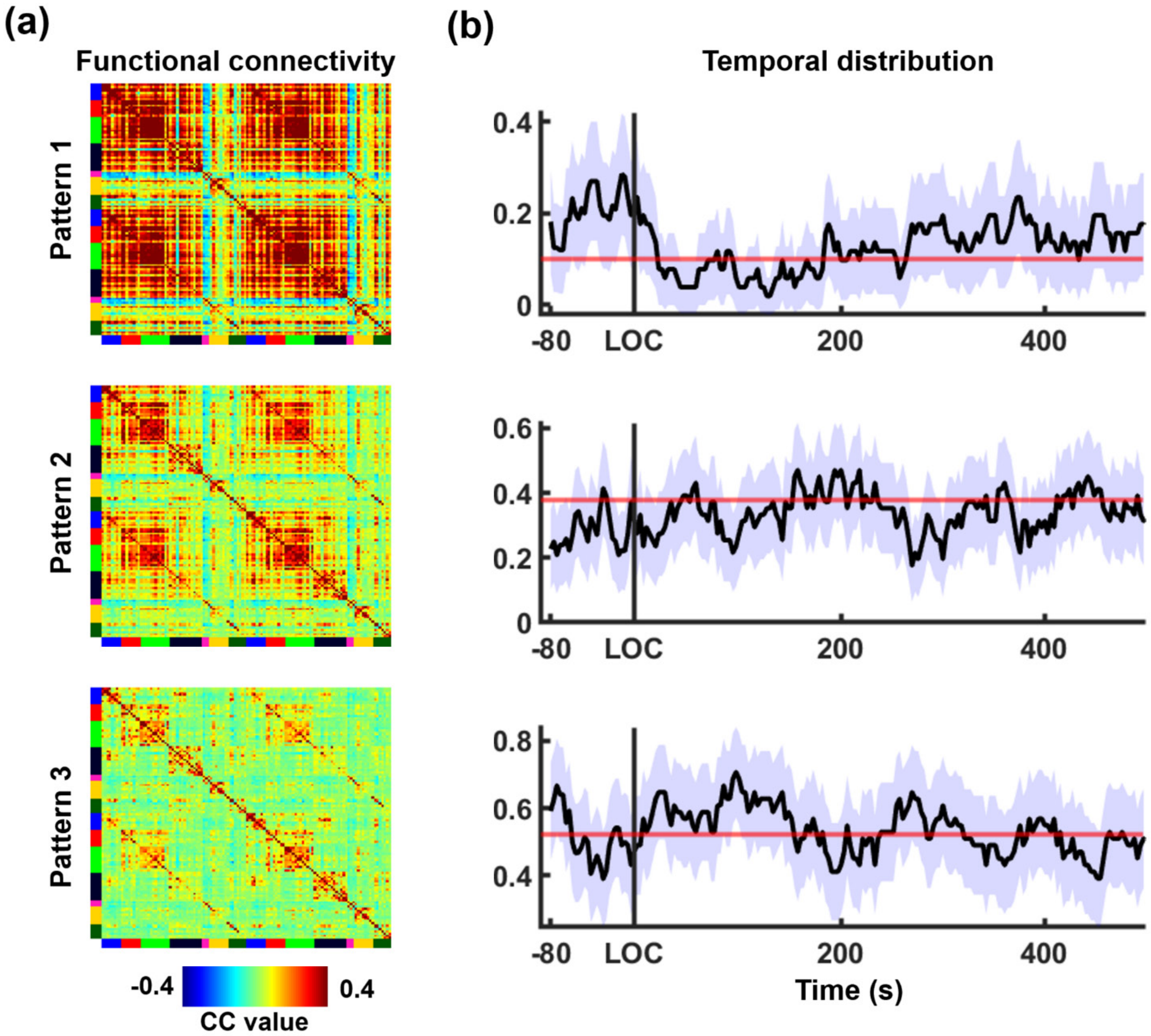


**Figure S11. Dynamic brain states under different consciousness conditions with a cluster number of 3.** (**a**) Three brain states, identified by unsupervised clustering of dynamic functional connectivity matrices. (**b**) Temporal distributions of three brain states around LOC and under the sustained unconsciousness. The purple shade indicates 95% confidence interval. The red line represents the averaged temporal distribution under low-dose propofol (i.e. baseline).

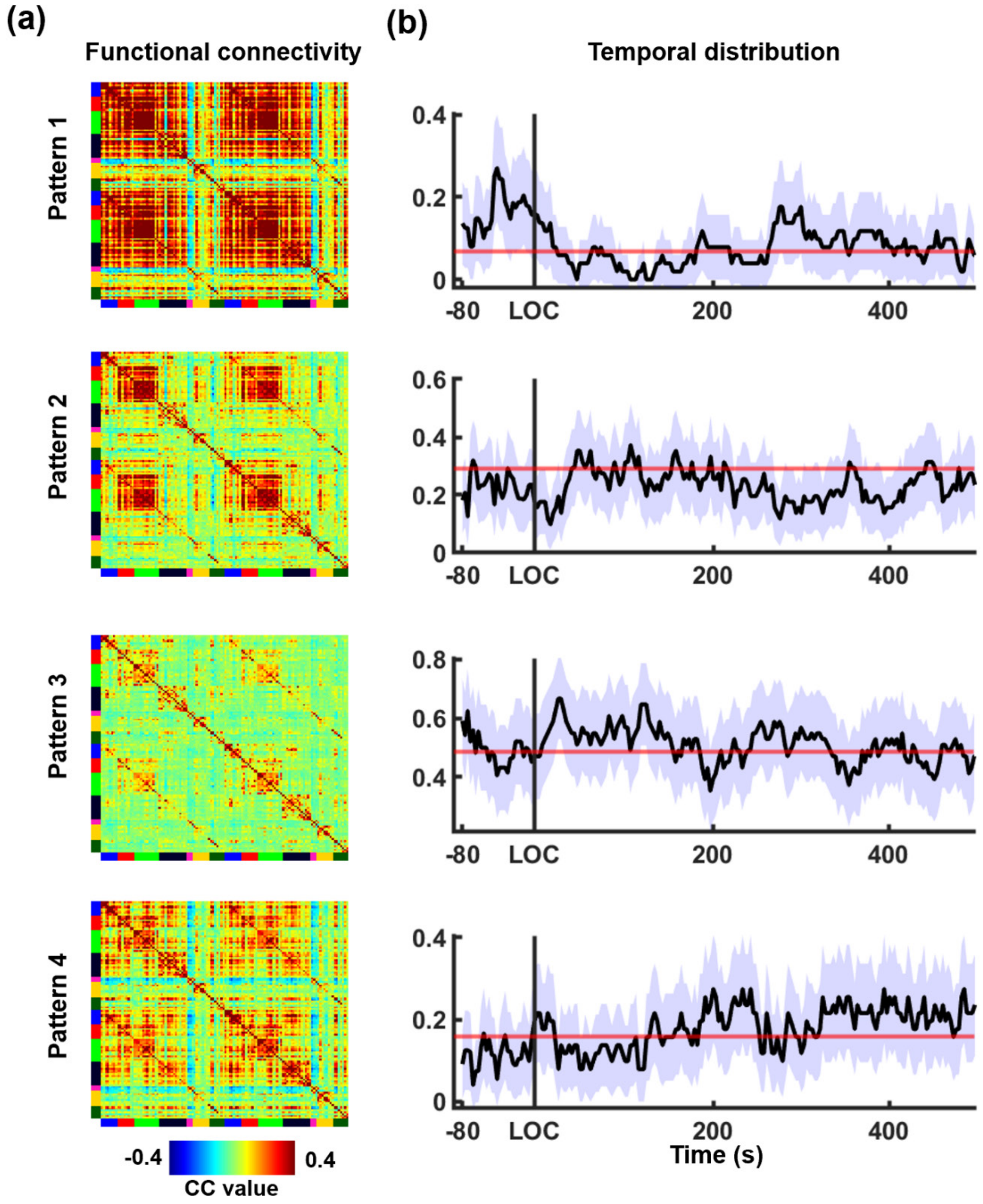


**Figure S12. Dynamic brain states under different consciousness conditions with a cluster number of 4.** (**a**) Four brain states, identified by unsupervised clustering of dynamic functional connectivity matrices. (**b**) Temporal distributions of four brain states around LOC and under the sustained unconsciousness. The purple shade indicates 95% confidence interval. The red line represents the averaged temporal distribution under low-dose propofol (i.e. baseline).

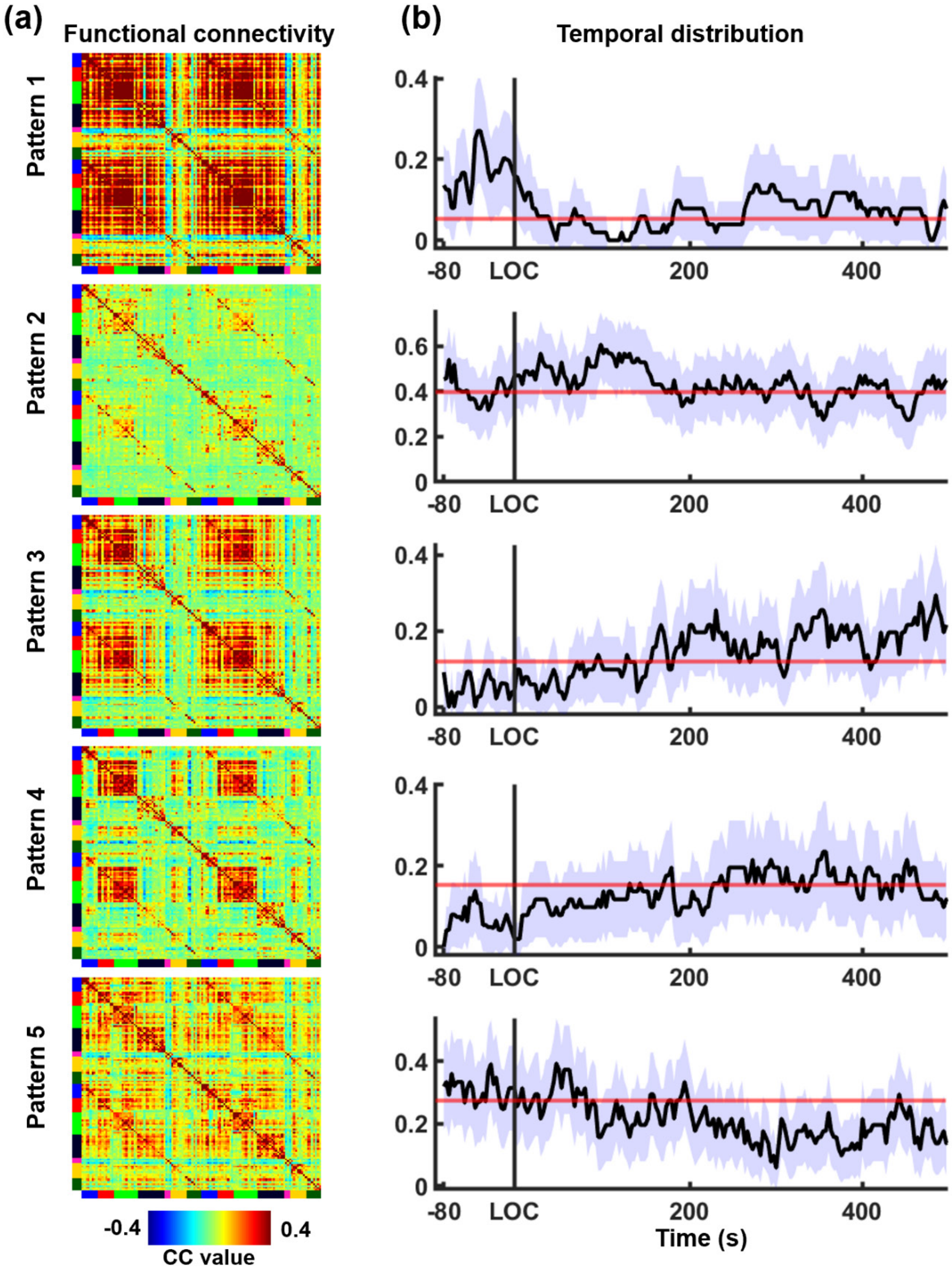


**Figure S13. Dynamic brain states under different consciousness conditions with cluster number of 5.** (**a**) Five brain states, identified by unsupervised clustering of dynamic functional connectivity matrices. (**b**) Temporal distributions of five brain states around LOC and under the sustained unconsciousness. The purple shade indicates 95% confidence interval. The red line represents the averaged temporal distribution under low-dose propofol (i.e. baseline).

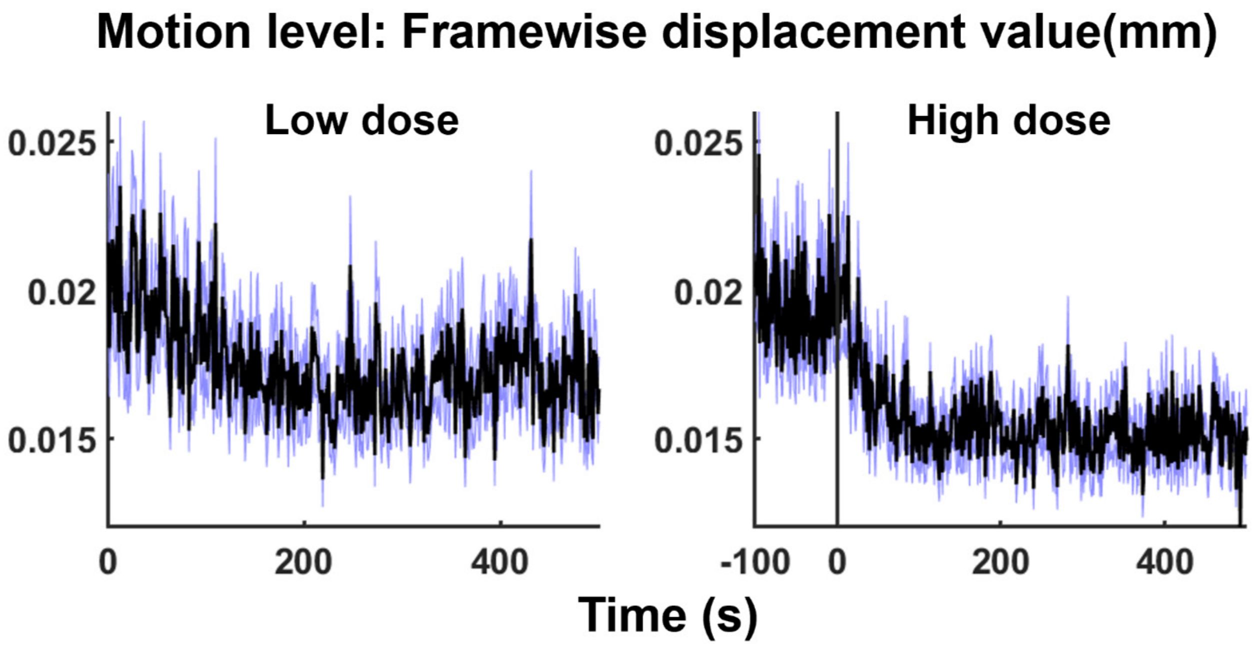


**Figure S14. Motion level under (a) low-dose propofol and (b) around loss of consciousness.** The moment of LOC is defined as time 0 for the second scan (i.e. high dose).

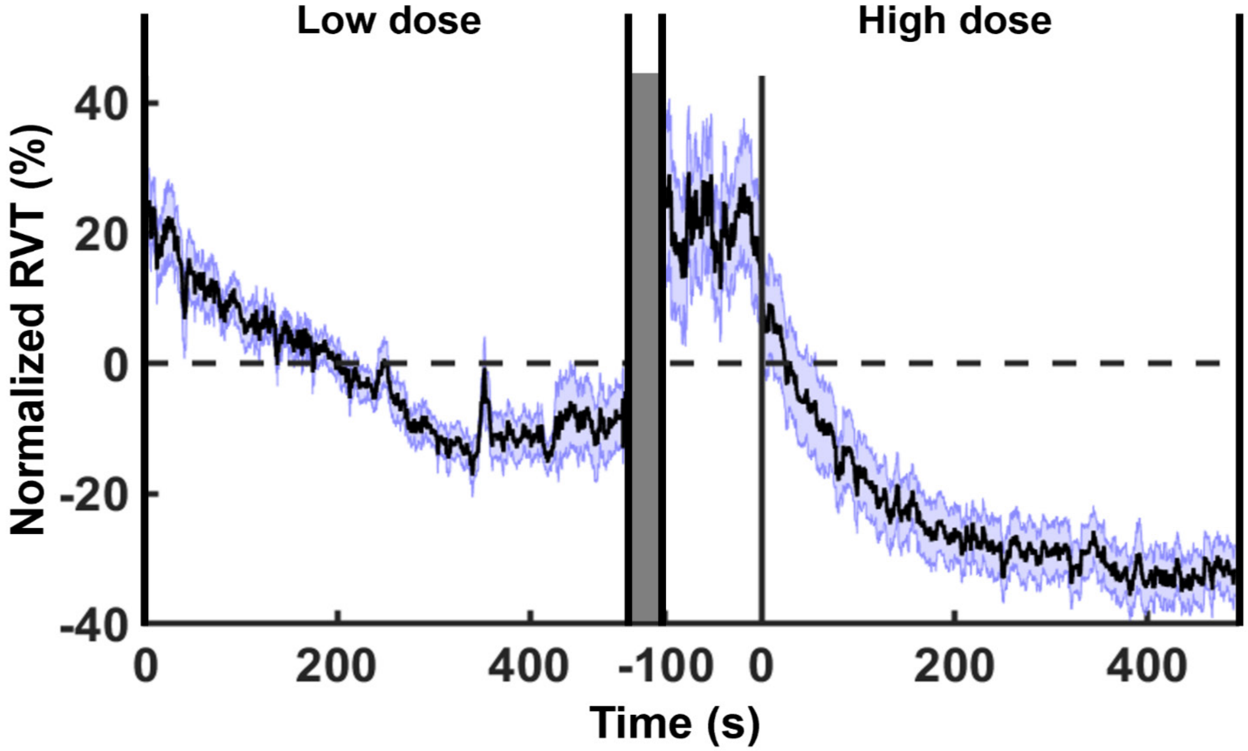


**Figure S15. Respiration during graded propofol.** The moment of LOC is defined as time 0 for the second scan (i.e. high dose).
